# Implantable Living Materials Autonomously Deliver Therapeutics from Contained Engineered Bacteria

**DOI:** 10.1101/2025.10.09.681377

**Authors:** Tetsuhiro Harimoto, Fernando Herrero Quevedo, Janis Zillig, Sanjay Schreiber, Yi Wu, Christine Heera Ahn, Tania To, Rohan Thakur, Alexander Tatara, Shawn Kang, Zheqi Chen, Shanda Lightbown, David Weitz, David J. Mooney

**Affiliations:** John A. Paulson School of Engineering and Applied Sciences, Harvard University, Cambridge, Massachusetts, USA; The Wyss Institute for Biologically Inspired Engineering, Harvard University, Boston, Massachusetts, USA; Harvard-Massachusetts Institute of Technology Division of Health Sciences and Technology, Massachusetts Institute of Technology, Cambridge, Massachusetts, USA; Division of Infectious Diseases and Geographic Medicine, The University of Texas Southwestern Medical Center, Dallas, Texas, USA; Department of Physics, Harvard University, Cambridge, Massachusetts, USA

## Abstract

Microbes are increasingly utilized as living therapeutic vehicles, yet their uncontrolled dissemination in the body has long remained a roadblock to clinical development. Physical containment, while widely used for mammalian cells, remains largely unattainable due to eventual bacteria escape. Here, we present an implantable material platform that encapsulates and confines bacteria, wherein synthetically engineered microbes produce therapeutic payloads from within. To prevent microbial escape, we developed a hydrogel scaffold with dual mechanical features: high stiffness to regulate bacterial proliferation and high toughness to resist material fracture under physiological stress. This design achieved complete bacterial containment for over six months and withstood multiple forms of mechanical loading that otherwise caused catastrophic material failure. By genetically engineering embedded bacteria, we endowed the material with environmental sensing and on-demand therapeutic release capabilities and demonstrated autonomous treatment in a murine prosthetic joint infection model. This multimodal strategy provides a safe and generalizable framework for deploying microbial medicines *in vivo* and supports their use as autonomous drug depots across a range of disease settings.

## Introduction

Synthetically engineered cells are emerging as living therapeutic modalities, capable of sensing physiological conditions and producing bioactive payloads *in vivo* (*1–5*). One promising class is bacteria that colonize a wide range of diseased and harsh physiological environments, including cytosolic space (*6–8*), mucosa (*9–12*), infected sites (*13–17*), epidermis (*18–21*), inflammatory tissues (*22–27*), and tumors (*28–31*). However, clinical success of microbial therapy is limited, largely due to unwanted dissemination of the microbes in the body that led to multiple clinical trial failures (*32–41*). While several genetic control strategies are being developed (*42–47*), evolutionary pressure often results in mutational escape.

Implantable hydrogels offer a physical strategy to confine therapeutic cells at target sites. Such living materials hold promise as localized drug depots with the capacity to dynamically respond to diseased environments (*48–50*). While previous efforts have largely focused on encapsulating mammalian cells, their inherent sensitivity to hypoxic and avascular implanted environments led to loss of viability (*51–53*). Given microbes’ robust viability in these conditions, bacterial encapsulation using implantable hydrogels presents an opportunity to bridge the complementarity of materials and living therapeutic vectors. Yet, vast previous work only reports short-term containment that often led to eventual bacterial escape from the encapsulating materials, and *in vivo* confinement within implants remains unrealized (*54–77*) (**table S1**).

We hypothesized that fulfilling two key criteria for a material enables robust and durable containment of therapeutic bacteria: (i) resistance to the internal forces generated by proliferating bacteria and (ii) mechanical toughness sufficient to withstand deformation from surrounding tissues. In addition, fabrication processes need to be biocompatible to preserve bacterial viability. Implantable Living Materials (ILMs) built on these principles achieve stable microbial containment *in vivo*, enabling disease sensing and autonomous therapeutic release (**Fig. 1A**).

**Fig. 1:**
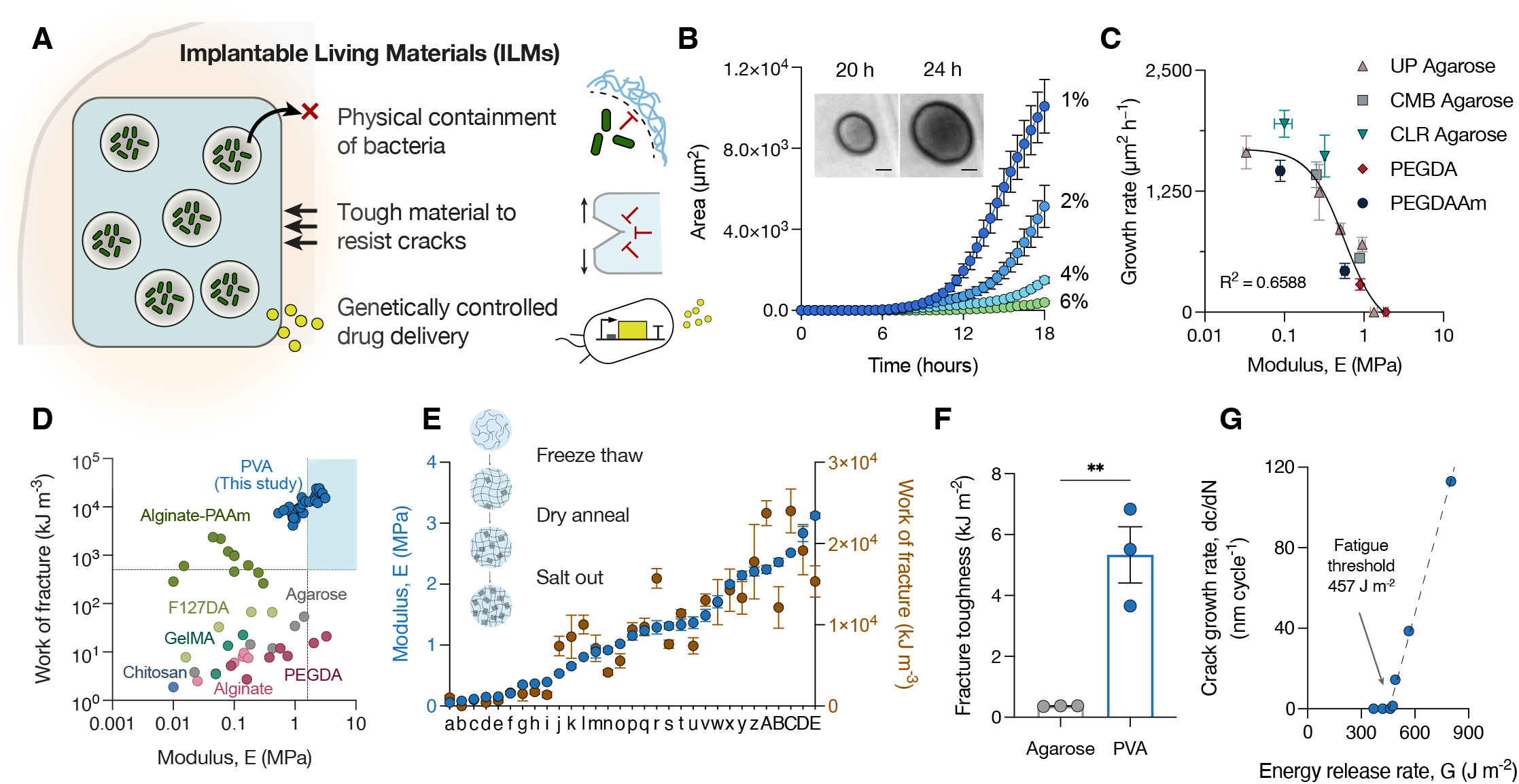
Design and mechanical characterization of Implantable Living Materials (ILMs). (A) Schematic of the ILM platform, in which a tough hydrogel scaffold encapsulates therapeutic bacteria. The material provides physical containment, resists both internal proliferative forces and external mechanical stresses, and enables genetically controlled therapeutic release. (B) Colony area of embedded *E. coli* ClearColi (Ecc) in agarose hydrogels of different concentrations over time. Insets: representative time-lapse microscopy images of colonies in 2% agarose (scale bar, 50 μm). Data are mean ± SEM; n ≥ 5 independent replicates. (C) Growth rate of encapsulated bacteria as a function of hydrogel stiffness. UP, UltraPure™ agarose; CMB, Certified Molecular Biology Agarose; CLR, Certified Low Range Ultra Agarose; PEGDA, Poly(ethylene glycol) diacrylate; PEGDAAm, Poly(ethylene glycol) diacrylamide. Data are mean ± SEM; n ≥ 2 independent replicates. (D) Ashby plot of hydrogel elastic modulus versus work of fracture. Blue points indicate the PVA hydrogel developed in this study; other data are from literature and in-house measurements (see Table S2). (E) Elastic modulus and work of fracture of PVA hydrogels fabricated under different processing conditions (see Table S3). Data are mean ± SEM; n ≥ 3 independent replicates. (F) Fracture toughness of agarose and PVA hydrogels with matched modulus (1.5 MPa), measured by pure shear testing. Data are mean ± SEM; all replicates are shown. *P < 0.05; unpaired *t*-test. (G) Cyclic crack growth of PVA hydrogel.

## Results

Bacteria generate mechanical forces through proliferation that are orders of magnitude greater than those produced by mammalian cells (*78–81*). To evaluate whether stiff encapsulating matrices can resist this biomass expansion, we fabricated biocompatible agarose hydrogels with increasing stiffness and tracked bacterial growth using time-lapse microscopy (**Fig. 1B; fig. S1, A and B; movie S1**). Colony expansion rates decreased as matrix stiffness increased (**Fig. 1B**), consistent with previous studies (*82, 83*), suggesting a mechanical feedback mechanism in which the surrounding material modulates bacterial growth. We next examined whether this relationship differs between bacterial strains used in therapeutic applications (*84–86*). Although the strains exhibited different baseline growth rates, all showed similarly reduced expansion as matrix stiffness increased (**fig. S2, A to C**). We selected *E. coli* ClearColi (Ecc) for subsequent studies, a strain commonly used in therapeutic contexts due to its high protein production and reduced immunogenicity (*87*).

To determine whether this stiffness-growth relationship holds across material platforms, we tested a library of hydrogels composed of different polymer types, concentrations, and molecular weights (**fig. S3, A to H**). Across all tested systems, higher stiffness consistently suppressed bacterial expansion (**Fig. 1C; fig S4, A to D**). Other mechanical properties, including tensile strength, work of fracture, and fracture strain, showed no consistent correlation with growth rate (**fig. S5, A to C**). Finally, we tested whether high stiffness could completely arrest Ecc growth. In both agarose and PEGDA systems, colony expansion ceased when matrix stiffness exceeded ~1.5 MPa (**Fig. 1C**). At this modulus, the bacteria were likely unable to deform the surrounding matrix, establishing a stiffness threshold required to constrain bacterial proliferation in ILMs.

Human tissues are subject to continuous movement, which imposes mechanical stress and can damage implanted materials (*88*). Thus, in addition to being stiff, ILMs must also be tough to prevent material fracture and bacteria escape *en masse*. However, increasing material stiffness often results in brittleness according to the Lake-Thomas theory (*89*). While several hydrogels have been engineered to overcome this trade-off, many rely on components and fabrication methods that are incompatible with living bacteria (*90, 91*). Previously reported hydrogels that encapsulated therapeutic *E. coli* did not attain sufficient stiffness and fracture toughness simultaneously (*92–95*) (**Fig. 1D, table S2**). As a benchmark for anatomical stress, we used reported work of fracture of soft collagenous tissue (*96*), ~0.5 MJ/m^3^.

To develop ILMs that are both stiff and tough, we turned to a class of hydrogels crosslinked through dense crystalline domains. This architecture of crystalline domains provide high stiffness by acting as rigid particles, while deconcentrating stress to resist crack growth (*97*). Polyvinyl alcohol (PVA) was selected as the base polymer due to its low toxicity, chemical stability, and established clinical use (*98*). Because fabrication processes that increase PVA crystallinity can compromise bacterial viability, we optimized three methods commonly used to induce crystallization: freeze-thaw cycles (*99, 100*), dry-annealing (*101, 102*), and the Hofmeister effect mediated salting-out process (*103, 104*). Freezing at −80 °C significantly reduced Ecc viability, but adjusting the temperature to −20 °C preserved bacterial survival while still promoting PVA crystallization (**fig. S6A**). Dry annealing typically requires complete desiccation at 100 °C, which is incompatible with Ecc. Lowering the annealing temperature to 37 °C and incorporating known desiccation protectants, alginate (*105*) or gelatin (*106*), substantially improved viability, even after 24 hours of drying (**fig. S6B)**. For the Hofmeister salting-out method, we tested three strong kosmotropic salts: sodium sulfate (Na_2_SO_4_), sodium citrate (Na_3_C_6_H_5_O_7_), and sodium phytate (Na_12_C_6_H_6_O_24_P_6_). While sodium sulfate had minimal effect on viability, sodium phytate and sodium citrate rapidly killed Ecc. Neutralizing the pH of the solution restored viability (**fig. S6, C and D**).

Using these optimized fabrication conditions (freeze-thaw cycles, dry-annealing, and salting-out), we next aimed to achieve sufficient stiffness and toughness in the PVA hydrogels for Ecc containment. We tested parameters including polymer molecular weight and concentration, freezing duration and temperature, number of freeze– thaw cycles, dry annealing time, and salt types and concentration (**table S3**). Several combinations produced hydrogels with elastic modulus of ~3 MPa and a work of fracture exceeding 20 MJ/m^3^ (**Fig. 1E; fig. S7**), surpassing the thresholds needed to resist bacterial expansion and withstand tissue-level mechanical stress. To more directly assess toughness, we measured the fracture toughness of PVA (table S3, v) and agarose hydrogels with matched stiffness using pure-shear testing (*93*). The resulting PVA hydrogels exhibited fracture toughness of > 5 kJ/m^2^, an order of magnitude higher than brittle agarose (**Fig. 1F; fig. S8 and S9**). Because implanted materials are subjected to repeated loading, we also evaluated fatigue resistance via cyclic crack growth testing (*91*). The toughened PVA showed a high fatigue threshold of ~450 J/m^2^, representing a >10-fold improvement over agarose (**Fig. 1G; fig. S9**). Other mechanical properties of PVA, including fracture strain and tensile strength, also exceeded those of agarose by at least an order of magnitude (**fig. S9**).

We next sought to establish ILMs by incorporating bacteria within the engineered PVA matrix. An ideal ILM should contain internal voids to support bacterial growth and activity. However, introducing such mechanically weak regions can compromise the bulk integrity of the scaffold. To balance these competing requirements, we calculated the fractocohesive length of bulk PVA to be ~400 μm, defined as the ratio of fracture toughness to work of fracture per unit volume. This length estimates the maximum defect size the material can tolerate without sacrificing mechanical performance (*107*). Based on this threshold, we fabricated sacrificial microgels to introduce Ecc into microscale voids within PVA (**Fig. 2A**). Specifically, Ecc-containing microgels were generated using a water-in-oil emulsion and filtered to remove particles exceeding the size limit (**fig. S10, A and B**). Gelatin was selected as the microgel material due to its ability to protect Ecc from desiccation and its thermoreversible gelation at physiological temperatures (**fig. S6B and S10C**). To determine the allowable microgel loading, we varied the volume fraction of microgels incorporated into the PVA matrix. Mechanical testing showed that bulk material properties were preserved at microgel fractions below ~20% (**fig. S11, A and B**), likely near the percolation limit (*108*).

**Fig. 2:**
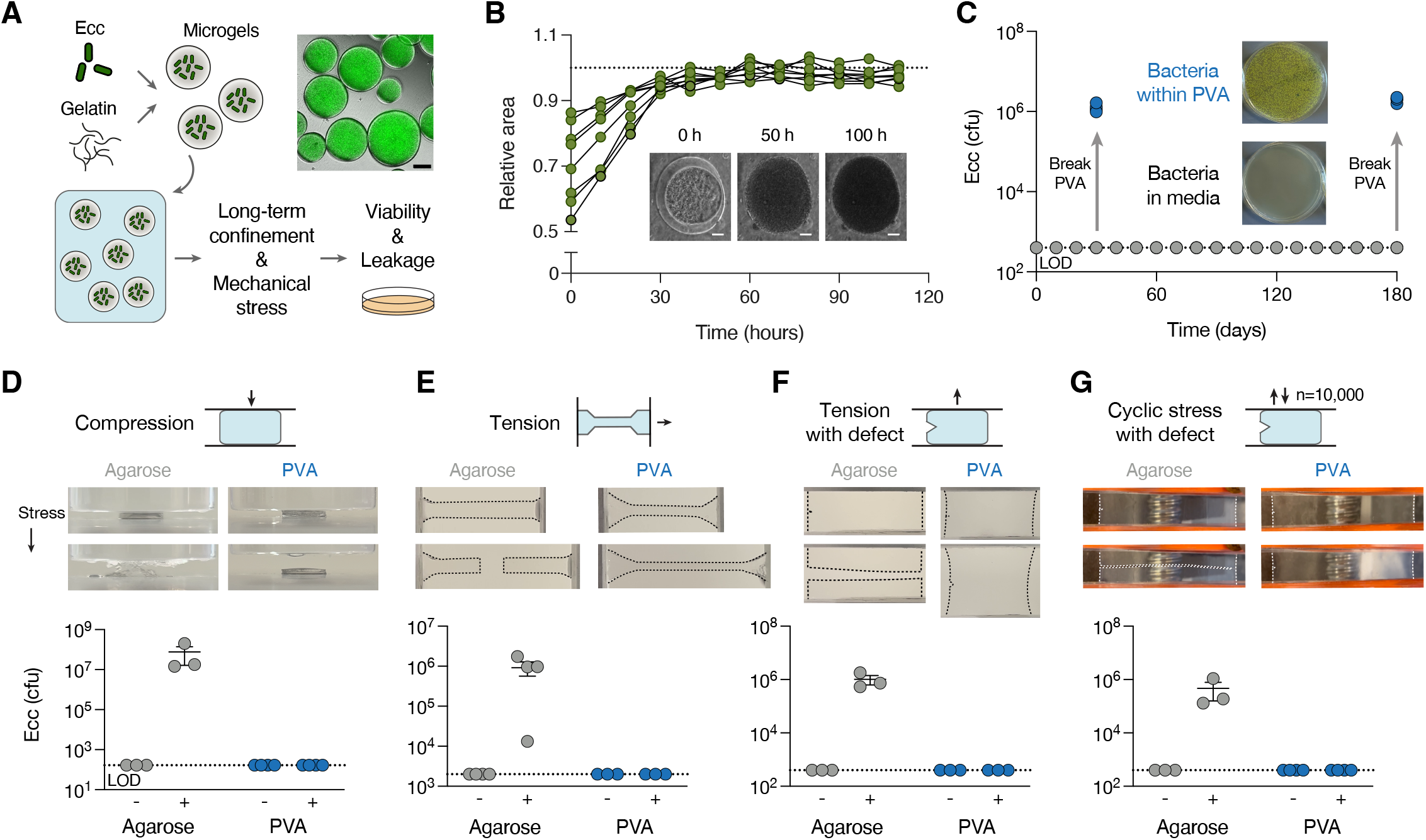
ILMs maintain bacterial containment during long-term culture and mechanical loading. (A) Fabrication of ILMs. *E. coli* ClearColi (Ecc) were encapsulated in sacrificial gelatin microgels by water-in-oil emulsion and subsequently embedded in a macroscale PVA hydrogel. Right: GFP-expressing Ecc in gelatin microgels (scale bar, 50 μm). (B) Colony expansion of Ecc within voids in the PVA matrix over time; dashed line indicates scaffold boundary. Insets: representative time-lapse microscopy images at indicated time points. (C) Long-term biocontainment of Ecc within ILMs. Samples were cultured for six months, with bacterial escape quantified every 10 days. Blue points indicate viable Ecc recovered from within the hydrogel; images show representative colonies on agar plates at day 30. Data are mean ± SEM; all replicates are shown. (D–G) Mechanical testing of PVA ILMs compared with agarose hydrogels under (D) compression, (E) tension, (F) tension with a notch defect, and (G) cyclic tensile loading (10,000 cycles). Defects in (F) and (G) simulated material flaws. Top: representative hydrogel images under load. Bottom: bacterial counts recovered before (−) and after (+) loading. Data are mean ± SEM; all replicates are shown. *LOD*, limit of detection; *cfu*, colony-forming units.

We tested whether ILMs could effectively prevent bacterial escape. Time-lapse microscopy of ILMs immersed in culture media showed initial Ecc replication within the microgel voids (**Fig. 2B**), confirming that bacterial viability was preserved during ILM fabrication. As colonies expanded, however, growth stalled upon reaching the interface between the microgels and the surrounding PVA matrix, with no observable invasion into the bulk scaffold. To assess long-term containment, ILMs were cultured continuously for up to six months. Intermittent sampling of the surrounding media revealed no detectable bacterial escape over this period (**Fig. 2C**). In parallel, bacteria within the PVA scaffold remained viable throughout the entire culture period.

To evaluate mechanical robustness, ILMs were subjected to a range of stresses and tested for bacterial leakage. Agarose hydrogels with matched stiffness served as controls. Under simulated compressive and tensile loading conditions (200-300 kPa for muscle contractions (*109*)), ILMs released no detectable Ecc, whereas agarose fractured and permitted bacterial escape (**Fig. 2, D and E; fig. S12, A and B; fig. S13, A and B; movie S2 and S3**). Post-loading viability assays confirmed that Ecc remained viable within intact ILMs after mechanical stresses (**fig. S12, A and B**). We also modeled potential material defects or damage by introducing a notch into the scaffold prior to tensile loading. While low stress levels around <150 kPa propagated cracks and released bacteria from agarose, ILMs resisted crack propagation and prevented escape even at ~500 kPa (**Fig. 2F; fig S12C; fig S13C; movie S4**). To simulate long-term exposure to physiological strain, ILMs were also subjected to cyclic tensile loading mimicking repeated physiological stress. After 10,000 loading cycles, the scaffold remained intact and continued to confine viable Ecc (**Fig. 2G; fig S12D; movie S5**).

One promising application of ILMs is their use as local drug depots. While implantable materials are widely used for local therapeutic delivery, conventional designs are limited in dosing dynamics, sustainability, and sensing capabilities (*110*). ILMs have the potential to overcome these limitations by genetically engineering embedded bacteria to sense disease and produce drugs on demand. As a proof-of-principle application, we focused on periprosthetic implant infections, which are often difficult to diagnose and would benefit from responsive therapeutic delivery to reduce the risk of dysbiosis and antibiotic resistance (*111*). Pseudomonas aeruginosa infection is particularly problematic due to its robust association with implant surfaces and inherent resistance to many common classes of antibiotics. Treatment failure can occur in up to ~75% of cases despite additional surgery, debridement, and intravenous antibiotic courses (*112*).

We designed a genetic circuit that enables autonomous sensing of P. aeruginosa, followed by on-demand therapeutic release through Ecc self-lysis (**Fig. 3A**). Specifically, a quorum-sensing system was constructed in Ecc by coupling the N-acyl homoserine lactone (AHL)-sensitive *luxB* promoter with the expression of a self-lysis *E* gene. In this system, bolus therapeutic release is achieved upon activation, while stochastic lysis enables microbial regeneration for subsequent activation (*113*). To evaluate sensing performance, we exposed Ecc harboring a GFP-coupled sensor circuit to purified AHL from *P. aeruginosa*. Quorum induction led to elevated GFP expression, with a limit of detection in the nanomolar range (**fig. S14A**). Next, we cloned the *E* gene downstream of the sensor which triggered Ecc lysis upon exposure to purified AHL and *P. aeruginosa* spent media, releasing GFP payload (**fig S14, B to E**). To enable antimicrobial activity, we built a therapeutic cassette expressing a synthetic anti-P. aeruginosa protein, chimeric pyocin (*114*) (ChPy), under a strong constitutive promoter. ChPy circumvents natural pyocin resistance found in certain Pseudomonas strains but also induces self-toxicity in Ecc. To mitigate this effect, we co-expressed multiple immunity genes in tandem, enabling stable ChPy production (**fig. S15**). To evaluate therapeutic efficacy, sonicated lysates from engineered Ecc were applied to P. aeruginosa cultures, resulting in a reduction in pathogen viability (**fig. S16, A and B**). Finally, integrating the sensor and therapeutic modules into a single strain enabled autonomous, AHL-triggered treatment of P. aeruginosa in multiple strains (**fig. S17, A and B**).

**Fig. 3:**
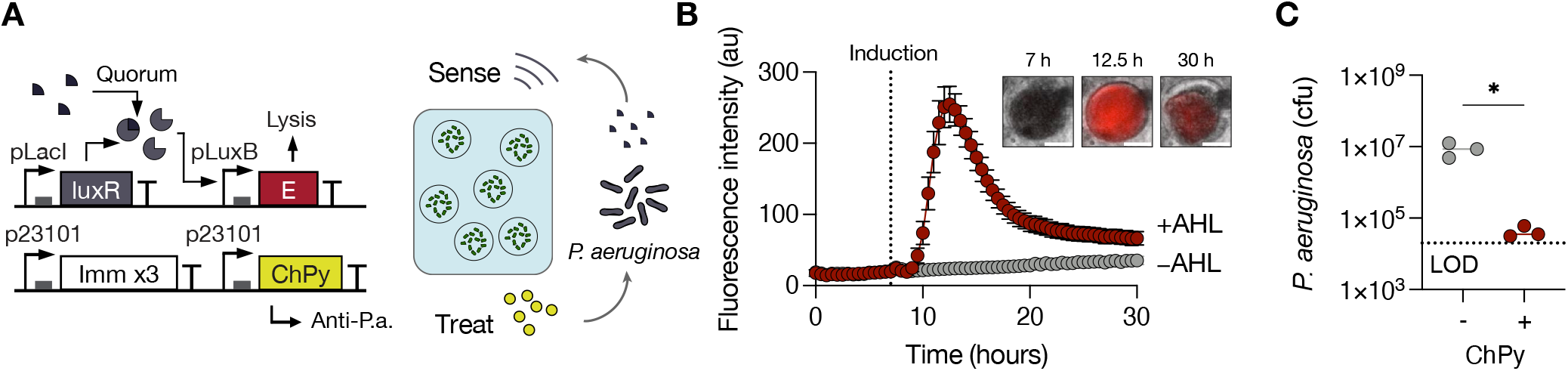
ILMs detect pathogen signals and release therapeutic payloads via engineered genetic circuits. (A) Genetic circuit design for pathogen sensing and eradication. *E. coli* ClearColi (Ecc) within ILMs constitutively expressed a synthetic anti–*P. aeruginosa* protein (chimeric pyocin, ChPy) together with three tandem immunity genes (Imm x3), and contained a *pluxB* promoter responsive to the quorum molecule acyl-homoserine lactone (AHL). Upon sensing quorum signals from *P. aeruginosa*, the circuit triggers self-lysis, releasing therapeutic payloads from ILMs. (B) Real-time sensing and lysis dynamics of embedded Ecc within ILMs. Fluorescence increased after AHL addition, followed by rapid loss due to lysis. Insets: representative time-lapse fluorescence microscopy images at indicated time points. Data are mean ± SEM; n ≥ 10 independent replicates. (C) *In vitro* efficacy of therapeutic ILMs. Reduction of *P. aeruginosa* was observed upon coculture with ILMs expressing ChPy. Data are mean ± SEM; all replicates are shown. *P < 0.05; unpaired *t*-test. *au*, arbitrary units; *LOD*, limit of detection; *cfu*, colony-forming units.

Having engineered the bacterial genetic circuit, we incorporated therapeutic Ecc into the PVA scaffold. We tracked the sense-and-respond behavior of the lysis circuit using time-lapse microscopy. Following AHL addition, fluorescent signal rose and then rapidly dissipated (**Fig. 3B; movie S6**), consistent with quorum activation followed by population collapse through programmed lysis. To assess long-term activity of the living material, we measured circuit activation after three and six months of culture. ILMs retained sensing functionality at this time point (**fig. S18, A and B**), demonstrating durable performance of the embedded genetic system. To evaluate the therapeutic release capability of ILMs, we measured diffusion of FITC-dextran molecules of varying sizes. The matrix exhibited a diffusion cutoff of ~250 kDa (**fig. S19A**), suggesting that many macromolecular protein payloads such as ChPy can be released over the course of days. Induced GFP expression from embedded Ecc confirmed protein production and release from within the ILM (**fig. S19B**). When cultured with *P. aeruginosa*, ILMs inhibited pathogen growth, demonstrating closed-loop infection treatment via autonomous sensing, lysis, and antimicrobial release (**Fig. 3C**).

We next evaluated the use of ILMs for periprosthetic joint infection *in vivo*. A murine model was established by implanting a stainless steel prosthesis into the femoral intramedullary canal (*115–118*), and tethering ILMs to the distal end of the implant to localize them within the joint space (**Fig. 4, A and B**). To assess bacteria containment, we measured Ecc dissemination into surrounding tissues following implantation. No Ecc was detected in tissues near the ILM after one day, whereas bacterial escape occurred in the absence of the PVA scaffold (**Fig. 4C**). We next assessed long-term safety and stability. Over the course of a week, mice gradually regained body weight (**fig. S20A**), consistent with typical post-surgical recovery (*119*). At days 1, 3, and 7 post-implantation, no detectable Ecc was found in periprosthetic tissue, the prosthetic device, liver, or spleen (**Fig. 4D; fig S20B**). Despite effective biocontainment, ILMs supported the growth of encapsulated Ecc to saturation and maintained their viability *in vivo*. To test whether the engineered genetic circuit remained functional after implantation, we isolated Ecc harboring the quorum-sensing plasmid from explanted ILMs. All recovered colonies retained the engineered plasmid (**fig. S21**). Upon AHL induction, all >100 tested isolates showed robust GFP expression (**Fig. 4E**), confirming that sensor function was preserved *in vivo*.

**Fig. 4:**
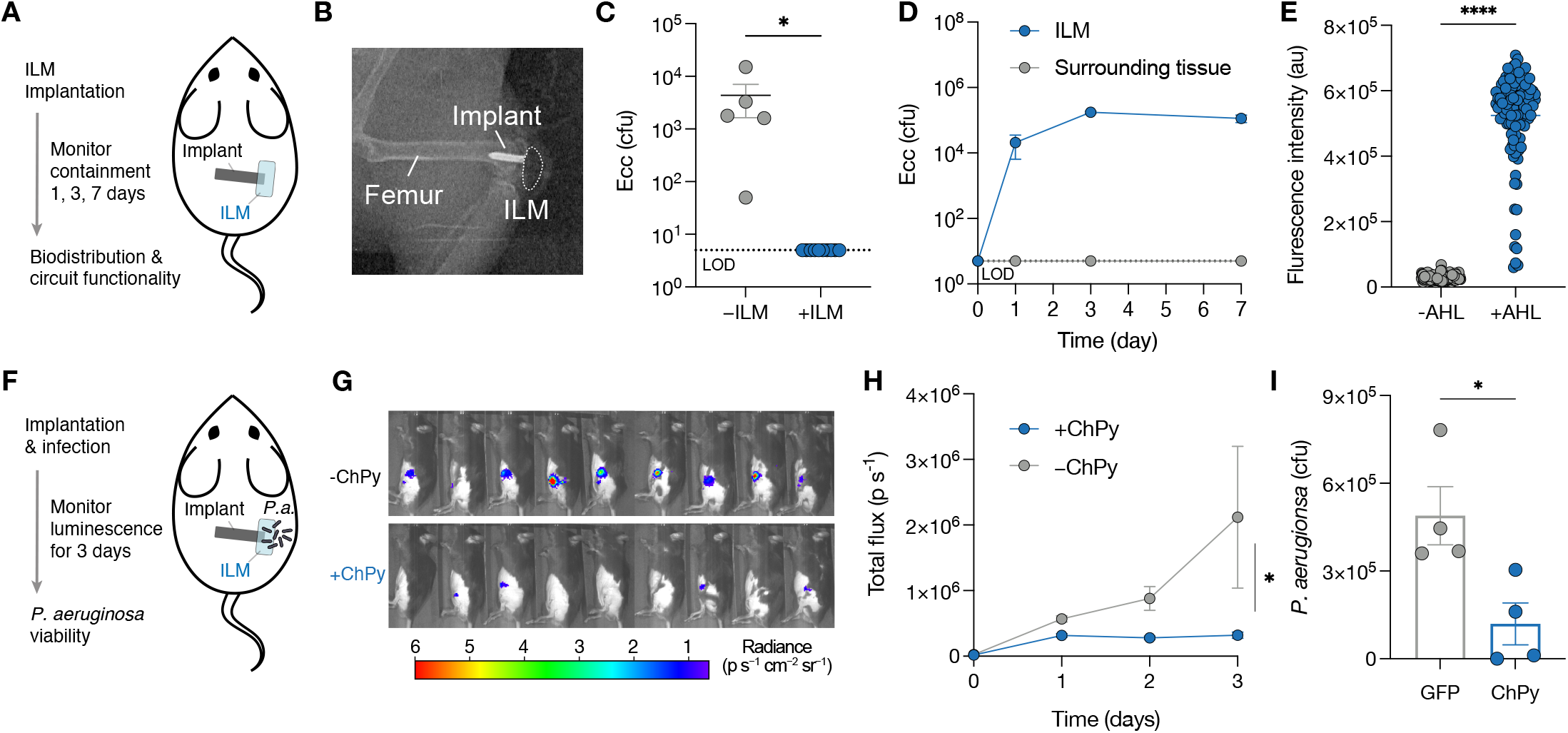
ILMs confine engineered bacteria and treat infection in a murine joint implant model. (A) Experimental design to assess *in vivo* containment and safety of ILMs. (B) Representative X-ray of a mouse femur implanted with a stainless-steel prosthesis and an ILM. (C) Viable *E. coli* ClearColi (Ecc) recovered from surrounding joint tissue one day after implantation. Data are mean ± SEM; all replicates are shown. *P < 0.05; unpaired *t*-test. (D) Bacterial counts in joint tissue and within ILMs over seven days. Data are mean ± SEM; n ≥ 5 independent replicates. (E) Genetic circuit function following implantation. Ecc were isolated from explanted ILMs and assayed for sensor response 24 hours after AHL induction. Data are mean ± SEM; all replicates are shown. ****P < 0.0001; paired *t*-test. (F) Experimental design for evaluating ILM efficacy against *P. aeruginosa* infection. (G) Representative IVIS images of bioluminescent *P. aeruginosa* at day 3 post-infection. (H) Quantification of *P. aeruginosa* bioluminescence over three days in mice treated with therapeutic ILMs (+ChPy) or control ILMs (−ChPy). Data are mean ± SEM; n ≥ 9 independent replicates. *P < 0.05; two-way ANOVA. (I) Viable *P. aeruginosa* (patient-derived strain) recovered from joint tissue at day 3. Data are mean ± SEM; all replicates are shown. *P < 0.05; unpaired *t*-test. *LOD*, limit of detection; *au*, arbitrary units; *cfu*, colony-forming units.

Finally, we tested the efficacy of ILMs by infecting the prosthetic device with *P. aeruginosa* (**Fig. 4F**). To track colonization of the pathogen in joint space, we used bioluminescent *P. aeruginosa* Xen41 (*118*). In mice treated with ILMs lacking ChPy, the luminescent signal continued to rise over three days (**Fig. 4, G and H**), indicating uncontrolled infection. In contrast, ILMs containing therapeutic Ecc suppressed *P. aeruginosa* growth, as seen by slower signal accumulation. To study ILM efficacy against an unmodified, clinically relevant pathogen, we next infected the mice with *P. aeruginosa* ATCC 15692 strain isolated from a patient’s wound (*120*). At three days post-infection, mice were sacrificed and viable *P. aeruginosa* levels at the implantation site were quantified. Compared to the control group, therapeutic ILMs significantly reduced the pathogen burden (**Fig. 4I**), demonstrating the ability of the system to autonomously sense and treat periprosthetic infection.

## Conclusion

We report the design of an implantable material platform that enables robust and long-term confinement of therapeutic bacteria in vivo. By engineering hydrogel mechanics beyond a critical stiffness and toughness threshold, the system prevents bacterial overgrowth and material fracture while maintaining cell viability over a long-term period. The fabrication process is fully compatible with living microbes, allowing incorporation of engineered strains without compromising material performance. Genetic programming of the encapsulated bacteria further equips the system with autonomous sense-and-respond capabilities, enabling on-demand therapeutic release in response to environmental cues. This integration of implantable scaffolds with living cellular functionality establishes a new class of dynamic drug delivery systems. While future efforts will focus on investigating host-device interactions and long-term effects (*121*), this platform lays the foundation for broader applications of engineered living materials in therapeutics, tissue regeneration, and immune modulation.

## Acknowledgments

We thank N. Archer for providing the Xen41 strain, and F. Jin for providing the sequence for ChPy. We thank members of the ImmunoMaterials group at the Wyss Institute for expert consultation with materials characterization. We thank T.C. Ferrante for his advice with microscope image acquisition and analysis. This work was supported by Harvard Materials Research Science and Engineering Center (DMR-2011754) and the Wyss Institute.

## Author contributions

T.H. and D.J.M. conceived the study. T.H., F.H.Q., J.Z., S.S., Y.W., C.H.A., T.T., R.T., and Z.C. designed and constructed material systems and performed all associated *in vitro* experiments. T.H., F.H.Q., and S.S. designed and constructed the genetic circuits and performed all associated *in vitro* experiments. T.H., F.H.Q., Y.W., A.T., S.K., and S.L. performed *in vivo* and/or *ex vivo* experiments. T.H., F.H.Q., and D.J.M. wrote and/or edited the manuscript with input from all authors. F.H.Q. and J.Z. contributed equally.

## Data and materials availability

All data are available in the main text or supplementary materials. Correspondence and request for materials should be addressed to D.J.M.

## Materials and Methods

### Hydrogel preparation

UltraPure™ Agarose (Thermo Fisher Scientific, 16500), Certified Low Range Ultra Agarose (Bio-Rad, 1613106), and Certified Molecular Biology Agarose (Bio-Rad, 1613100) were dissolved in Milli-Q water at the indicated concentrations. Solutions were microwaved with intermittent stirring until fully dissolved. To minimize evaporation, beakers were covered with wetted Kimwipes during heating. Molten agarose was cast between two glass plates separated by spacers and allowed to cool to room temperature for thermal gelation. Poly(ethylene glycol) diacrylate (PEGDA; MW 700 Da; Sigma-Aldrich, 455008) and poly(ethylene glycol) diacrylamide (PEGDAAm; MW 3700 Da; Sigma-Aldrich, 725676) were dissolved in Milli-Q water and dissolved by mixing. For photopolymerization, lithium phenyl-2,4,6-trimethylbenzoylphosphinate (LAP; Sigma-Aldrich, 900889) was added at 5 mg mL^−1^. Precursors were cast between a glass slide and a 0.18 mm coverslip with spacers and exposed to 365 nm UV light (36 W NailStar lamp) for 3 minutes. Hydrogels composed of Pluronic F127 (Plu) and Pluronic F127 diacrylate (PluDA) were prepared as previously described(*67*). Briefly, Pluronic F127 (Sigma-Aldrich, SLBG6026V) and Pluronic F127-DA (Creative PEG-Works, PPO-121) were dissolved in Milli-Q water to 30% (w/v) and mixed at the desired ratio. To prevent premature gelation, all steps were performed at 4 °C. Precursor solutions were cast and allowed to undergo physical gelation at room temperature for 10 minutes, followed by UV polymerization at 365 nm for 3 minutes. Polyvinyl alcohol (PVA; MW 89,000–98,000 Da, 146,000–186,000 Da, or 30,000–70,000 Da; Sigma-Aldrich, 341584, 363065, or 8136) solutions were prepared by microwave heating. Solutions were centrifuged at 4,000 × g for 2 minutes to remove air bubbles. For freeze–thaw gelation, PVA solutions were cast between glass slides separated by spacers, clamped, frozen, and thawed at room temperature. Hydrogels were dried at 37 °C for ~2 hours to remove water, immersed in pH-neutralized kosmotropic salt solutions for 24 hours, and then incubated in PBS for 24 hours to reach equilibrium swelling.

### Bacterial strains and culture conditions

*Escherichia coli* (ClearColi BL21(DE3), DH5α, and Nissle 1917), *Pseudomonas aeruginosa* (ATCC BAA-47, ATCC 15692, and Revvity Xen41), and *Bacillus subtilis* (ATCC 3610) were cultured in LB broth (Research Products International, L24065) supplemented with antibiotics as appropriate: 100 μg mL^-1^ ampicillin, 50 μg mL^-1^ kanamycin, 50 μg mL^-1^ spectinomycin, and 10 μg mL^-1^ tetracycline. Cultures were incubated at 37 °C with shaking at 250 rpm (New Brunswick Scientific). For lysis circuit strains, 0.2% glucose was added to overnight cultures to minimize leaky expression from the LuxR-regulated promoter.

### Bacteria growth within hydrogels

Overnight bacterial cultures were diluted into hydrogel precursor solutions, which were cast to 1 mm thickness and crosslinked using polymer-specific protocols. Hydrogel disks (2 mm diameter) were generated with a biopsy punch (Sklar, 06-4142). Each disk was placed into a 48-well glassbottom plate (MatTek, P48G-1.5-6-F) and secured within 80 μL molten agarose. Wells were filled with 1 mL LB medium supplemented with appropriate antibiotics. Brightfield and fluorescence time-lapse imaging was performed, and growth rates were calculated using the image processing pipeline described below.

### Microscopy and image processing

Bacterial growth and ILM circuit dynamics were monitored by multipoint time-lapse microscopy. Imaging was performed at 37 °C on a Nikon Eclipse Ti2 microscope equipped with an Okolab incubation enclosure and a Zyla sCMOS camera (Andor Technology), controlled by Nikon Elements software. Images were acquired every 30 minutes in brightfield, GFP (Ex/Em 488/535 nm), and/or Cy5 (Ex/Em 640/670 nm) channels. Movies were analyzed using ImageJ (v1.53k). Frames were registered using the built-in SIFT algorithm prior to analysis. To monitor confined bacterial growth, individual colonies were background-subtracted and segmented in each frame using Otsu’s thresholding method. Growth rates were calculated by linear regression of colony area over time. To monitor ILM circuit dynamics, regions of interest (ROIs) were manually defined, and pixel intensities within them were tracked over time.

### Microgel preparation

A 30% (w/v) gelatin solution (Sigma-Aldrich, G2625) was prepared at 42 °C. Overnight *E. coli* cultures (OD_600_ = 3.5) were pelleted by centrifugation (3,000 × g, 10 minutes, 4 °C) and resuspended in gelatin at 50-fold concentration. This suspension was transferred into a 50 mL conical tube containing twice the volume of prewarmed 2% (w/w) surfactant (008-FluoroSurfactant, RAN Biotechnologies) in HFE-7500 (3M Novec). The mixture was vortexed at 2700 rpm for 60 seconds to generate a water-in-oil emulsion. Gelatin droplets were crosslinked by placing the tube on ice. Following gelation, microgels were washed three times with HFE oil to remove excess surfactant, demulsified using an antistatic gun (Milty Pro Zerostat 3), and washed three times with PBS to remove residual oil. Microgels <300 μm were recovered by filtering through a 300 μm strainer (pluriSelect) and stored at 4 °C.

### ILM fabrication

Bacteria–gelatin microgels were mixed into PVA solution (final microgel volume fraction, 20% unless otherwise noted). The mixture was either cast between glass slides separated by spacers and clamped, or molded in multiwell plates. Samples were frozen for 1 hour and thawed at room temperature, then dip-coated in fresh PVA solution and subjected to a second freeze–thaw cycle. Hydrogels were dried at 37 °C for ~2 hours to remove water, immersed in pH-neutralized kosmotropic salt solutions for 24 hours, and then incubated in PBS for 24 hours to reach equilibrium swelling. For *in vivo* studies, ILMs were fabricated as thin hemispherical shells anchored to stainless steel pin implants. To mold and tether the ILMs, 10 μL of PVA/bacteria–microgel precursor mixture was dispensed into the bottom of 1.5 mL tubes. Bacteria-free PVA was extruded through a 25-gauge blunt-end needle and inserted into the precursor solution. The needle was secured in place by piercing it through a plastic insert (filter from a trimmed P1000 pipette tip) positioned within the tube. Constructs were crosslinked as described above, and the distal ends of the pins were trimmed to ~0.7 cm using sterile wire cutters.

### Bacteria viability assays

To assess viability after freeze–thaw cycles, 1 mL of overnight *E. coli* culture was suspended in PBS and subjected to repeated freezing at −20 °C or −80 °C. To evaluate viability after drying, bacteria were mixed with 100 μL PBS, gelatin (10% w/v), or alginate (5% w/v), dried at 37 °C for 2 hours, and rehydrated in 1 mL PBS. For salting-out assays, bacterial cultures were incubated for 24 hours at room temperature in sodium sulfate (0.1–1.8 M), sodium citrate (0.1–1.8 M, with or without pH neutralization), or sodium phytate (0.1–0.3 M, with or without pH neutralization). For all assays, viability was determined by serial dilution and plating on agar to enumerate colony-forming units (CFU).

### Uniaxial tensile test

Tensile tests were performed using a TA.XT.plus Texture Analyzer (Stable Micro Systems) equipped with a 5 kg load cell. Dog-bone-shaped specimens were cut using an ISO 527-2-5B cutting die. For testing, each sample was clamped and stretched until rupture. The strain rate is fixed at 1 mm s^-1^ unless stated otherwise. Nominal stress (*σ*) was calculated by dividing recorded force (*F*) by cross-sectional area of undeformed state (*A*_0_). Strain (*ε*) was calculated as *ε* = (*L* − *L*_0_)/*L*_0_, where *L* is deformed length and *L*_0_ is initial length. The elastic modulus (*E*) was estimated from the slope of the stress-strain curves in linear elastic region which is up to 5% strain. The work of fracture (*W*_c_) was calculated by numerically integrating the area under the stress– strain curve up to the fracture strain 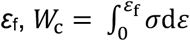.

### Fracture toughness measurement

Fracture toughness was determined as previously described using a pure shear method(*122, 123*). A sample of rectangular hydrogel with a width of 50 mm and height of 30 mm was prepared. Microscope slides were adhered to the long edges using cyanoacrylate glue (Krazy Glue) to serve as grippers. The stretchable region of the sample is 50 mm in width and 10 mm in height. Two samples were prepared in the same geometry. One sample was cut by a fresh razor blade to form a 20 mm precrack, and the other one remained intact. When measuring toughness, the intact sample was stretched to obtain a stress-strain curve *σ*(*ε*). The strain energy density was calculated as a function of strain, 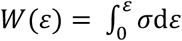. The precracked sample was stretched to fracture to obtain the critical strain, *ε*_c_. Fracture toughness (*G*_c_) was calculated as *G*_c_ = *W*(*ε*_c_)*H*, where *H* is the undeformed height of the precracked sample.

### Fatigue measurement

Fatigue tests used the same sample geometry as the toughness measurement. For the fatigue test, the strain energy density, *W*(*ε*), was measured from the stress–strain curve of the intact sample after 1,000 cycles of loading and unloading, such that the stress-strain curve reached a steady state. The precracked sample was applied to a cyclic strain (*ε*) of fixed amplitude. An amplitude of the strain corresponds to an amplitude of energy release rate, *G* = *W*(*ε*)*H*, where *H* is the undeformed height of the precracked sample. The crack growth was measured by an optical microscope after 10,000 cycles and divided by the number of cycles to get the crack growth per cycle (d*c*/d*N*). The microscope has a resolution of 5 μm, so that the resolution of d*c*/d*N* in our experiment is 0.5 nm/cycle. The loading rate in the fatigue test is 1 Hz. dc/d*N* was plotted against the amplitude of energy release rate *G*. The linear regression of data estimates the fatigue threshold (*G*_th_). G_th_ for agarose was roughly estimated due to its brittleness.

### Fractocohesive length

The fractcohesive length (*L*_f_) is calculated as the ratio of the fracture toughness (*G*_c_, in J/m^2^) to the work of fracture (*W*_c_, in J/m^3^), *L*_f_ = *G*_c_/*W*_c_. This length scale represents the critical scale of flaw size below which the material is insensitive to the flaw. For example, the strength and the stretchability of the hydrogel decrease in the presence of flaws larger than *L*_f_, and remain unaffected by flaws smaller than *L*_f_.

### Bacterial growth and confinement within PVA *in vitro*

Bacterial growth and confinement within ILM voids were monitored by time-lapse microscopy. ILMs were immobilized in glass-bottom plates (MatTek, P24G-1.5-10-F) by embedding in 2% (w/v) agarose, and each well was filled with 2 mL LB medium supplemented with appropriate antibiotics. Time-lapse imaging was performed every 30 minutes for 5 days at 37 °C, with medium refreshed every 24 hours. Quantification of bacterial growth within microgels was performed using the image processing pipeline described above. For long-term confinement and viability assays, ILMs were cultured in medium that was replaced every 7 days. To monitor bacterial escape, 5 μL of culture medium was sampled every 10 days. To quantify viability of encapsulated bacteria, PVA hydrogels were fragmented with a sterile razor blade, resuspended in PBS, and plated to enumerate CFU.

### Bacterial confinement under mechanical stress

All tools and instruments were sterilized with 70% ethanol prior to testing. Tensile tests were performed at 0.1 mm s^-1^ until reaching 500 g. Compression tests were performed at 0.1 mm s^-1^ until reaching 4,000 g. Tensile tests with defects were conducted on hydrogels containing ~1 mm cracks at 0.1 mm s^-1^ until reaching 2,000 g. Cyclic tensile loading was carried out at 1 Hz for 10,000 cycles to 7.5% strain. After loading, damaged regions of each sample were immersed in sterile PBS, and 5 μL of the solution was plated on agar to quantify CFU. To confirm bacterial viability within PVA, gels were manually fragmented to release encapsulated bacteria, resuspended in sterile PBS, and plated on agar to quantify CFU.

### Plasmid construction and gene circuit design

Plasmid pAJM.474 (#108526) was obtained from Addgene(*124*). All other constructs were assembled using Gibson Assembly. Plasmid backbones were amplified by PCR from existing plasmids, and genes of interest were obtained either as gBlocks (Integrated DNA Technologies) or by PCR. Constructs were transformed into *E. coli* DH5α or ClearColi™ BL21(DE3) and verified by longread Oxford Nanopore sequencing (Plasmidsaurus). Promoters, ribosome binding sites (RBSs), and circuit architecture were tuned to modulate expression. A complete list of plasmids is provided in Supplementary Table S4.

### Sensor and lysis circuit characterization

*P. aeruginosa* (ATCC 10145) was grown overnight in LB. Cultures were centrifuged, and the supernatant was filtered through 0.22 μm Steriflip™ filters (MilliporeSigma) and stored at 4 °C until use. *E. coli* strains harboring quorum-sensing (Plasmid #1, Table S4) and/or lysis (Plasmids #2 or #3) circuits were cultured overnight in LB with appropriate antibiotics, diluted 1:100 into fresh LB with antibiotics, and transferred to 96-well plates (Genesee Scientific). Cultures were incubated at 37 °C with shaking in a plate reader (BioTek Synergy H1) until reaching OD_600_ = 0.3–0.4, then treated with exogenous 3OC12-HSL (10 μM to 100 pM; Sigma-Aldrich) and monitored for 12–16 hours. Alternatively, cultures were treated with *P. aeruginosa* culture supernatant (2-fold or 10-fold dilution). GFP fluorescence was recorded every 20 minutes. Sensor and lysis circuit dynamics within ILMs were characterized by time-lapse microscopy. Dip-coated ILM disks were placed into glass-bottom plates (MatTek, P24G-1.5-10-F), embedded in 2% agarose, and incubated in LB at 37 °C. After 12–16 hours, cultures were induced with 100 μM 3OC6-HSL (Sigma-Aldrich). Brightfield and fluorescence images were acquired every 30 minutes for ~24 hours.

### Chimeric pyocin efficacy

*E. coli* strains expressing chimeric pyocin (ChPy) were grown overnight in LB with antibiotics, diluted 1:100 into fresh LB, and cultured to OD_600_ = 0.3–0.4. Cultures were centrifuged at 3,000 × g for 10 minutes and washed twice in sterile PBS. Pellets were resuspended in antibiotic-free LB, normalized by OD_600_, and lysed either by sonication (50% amplitude, 10 seconds on/45 seconds off, for a total time of 8 minutes) or by induction with 100 μM 3OC6-HSL (Sigma-Aldrich) for 3 hours at 37 °C. Lysates were filtered through 0.22 μm Steriflip™ filters and stored at 4 °C until use. Efficacy against *P. aeruginosa* was evaluated using spot and liquid culture assays. For spot assays, *P. aeruginosa* was spread on LB agar plates and allowed to dry before spotting with 5 μL of ChPy lysate. Plates were incubated overnight at 37 °C and examined the following day for zones of clearance. For growth inhibition in liquid culture, ~10 CFU of freshly grown *P. aeruginosa* were inoculated into 96-well plates with or without ChPy lysate. Cultures were incubated at 37 °C with shaking for 12–16 hours in a plate reader (Agilent Technologies), and OD_600_ was measured every 20 minutes.

### Drug release assay

Diffusion of fluorescein isothiocyanate–dextran (FITC–dextran) from PVA hydrogels was assessed using four molecular weight variants: 3,000–5,000 Da (FD4), 40,000 Da (FD40S), 250,000 Da (FD250S), and 2,000,000 Da (FD2000S) (all Sigma-Aldrich). Each was dissolved in PVA at 1 mg/mL, mixed thoroughly, and cast to a thickness of 1 mm between glass slides. PVA hydrogels were fabricated as described above. During salting-out and PBS swelling, hydrogels were incubated in PBS containing 1 mg/mL of the corresponding FITC–dextran to minimize premature leaching. Samples were rinsed briefly in fresh PBS and transferred to 24-well plates containing 500 μL PBS. At 2, 4, 6, 8, and 24 hours, 200 μL of PBS were collected and transferred to a 96-well plate for fluorescence quantification (λ_ex = 480 nm, λ_em = 530 nm) using a plate reader (BioTek Synergy H1). After each measurement, samples were placed into fresh PBS for continued release. A standard curve was generated from serial dilutions of 1 mg/mL FITC–dextran. Cumulative release (%) was calculated by normalizing the sum of detected mass across time points to the theoretical total FITC–dextran initially present in the hydrogel. GFP release from ILMs was assessed by encapsulating *E. coli* carrying sensor (Plasmid #1, Table S4), lysis (Plasmid #3), and GFP (Plasmid #5) circuits. ILMs were cultured in LB with 0.2% glucose for 8 hours, then transferred to LB without glucose and incubated overnight at 4 °C. ILMs were induced with 10 μM 3OC6-HSL (Sigma-Aldrich), and released GFP was quantified by sampling the culture medium and measuring fluorescence using a plate reader.

### ILM-*P. aeruginosa* coculture

ILMs were pre-cultured for 6 hours in LB supplemented with appropriate antibiotics and 0.2% glucose. For coculture, ILMs were transferred to 50 mL conical tubes, and ~10 CFU of *P. aeruginosa* were added directly on top. Tubes were capped to prevent dehydration and incubated at 37 °C for ~18 hours. After incubation, ILMs were resuspended in 0.5 mL PBS, mixed thoroughly by pipetting, and vortexed at 2700 rpm for 60 seconds. The suspension was plated on agar to enumerate *P. aeruginosa* CFU.

### *In vivo* model of periprosthetic joint infection

All animal experiments were approved by the Institutional Animal Care and Use Committee at Harvard University (protocol 22-01-408). Male C57BL/6 mice (9–16 weeks old; The Jackson Laboratory) were used. A model of *P. aeruginosa* periprosthetic joint infection (PJI) was established as previously described(*115–118*). Briefly, a small medial incision was made over the left knee of anesthetized mice (isoflurane: 3–4% for induction, 1–2% for maintenance) to expose the quadriceps–patellar complex. The complex was laterally displaced to expose the distal femoral head. The femoral intramedullary canal was manually reamed through the intercondylar notch using a 25-gauge needle (Becton Dickinson), followed by reaming with a 23-gauge needle to widen the hole. Before needle removal, a radiograph was taken to confirm canal placement and ensure the femur had not fractured. A stainless steel pin (~0.7 cm length, 0.52 mm diameter) with an ILM tethered to its distal end was then surgically inserted into the canal, positioning the ILM within the periprosthetic joint space. The implant was inoculated with ~100 CFU of *P. aeruginosa*. The quadriceps– patellar complex was repositioned, and the incision was sutured with 5-0 Prolene (Ethicon). Buprenorphine (0.05–0.1 mg kg^-1^) was administered subcutaneously as an analgesic, given preemptively and postoperatively every 12 hours for 72 hours.

### *In vivo* bacterial confinement

Implantable ILMs containing Ecc engineered with inducible GFP (Plasmid #6, Table S4) were fabricated as described above. Mice underwent surgical implantation of ILMs (“+PVA” group) or non-encapsulated microgels (“–PVA” group), without *P. aeruginosa* inoculation. Body weight was monitored daily. To assess bacterial escape and dissemination, mice were euthanized 1, 3, or 7 days post-implantation. Organs (spleen and liver), surrounding joint tissues, and metal implants were collected in bead-beating tubes (2.4 mm metal beads; Revvity) and mechanically homogenized using a Retsch MM 400 tissue dissociator (30 Hz, 3 minutes). ILMs from the “+PVA” group were also harvested, sectioned, and homogenized to quantify encapsulated bacterial content. Homogenates were serially diluted in sterile PBS and plated on LB agar with or without antibiotics. Colonies were counted after overnight incubation at 37 °C. Spleens and livers were weighed prior to dissociation to normalize recovered CFU counts.

### *In vivo* ILM efficacy

ILMs containing Ecc engineered with quorum-sensing (Plasmid #1, Table S4), lysis (Plasmid #3), and payload circuits (Plasmid #4 or #5) were fabricated, cultured *in vitro* for 12–16 hours, and surgically implanted into mice. *P. aeruginosa* strains were inoculated at the time of surgery as described above. Body weight was monitored daily. For the Xen41 strain, bioluminescence was measured daily using an IVIS Spectrum imaging system (PerkinElmer) and quantified with Living Image software. For ATCC 15692 strain, mice were euthanized 3 days post-implantation. Surrounding joint tissues, metal implants, and ILMs were collected in beadbeating tubes (2.4 mm metal beads; Revvity) and mechanically homogenized using a Retsch MM 400 tissue dissociator (30 Hz, 3 minutes). Homogenates were serially diluted in sterile PBS and plated on LB agar. *P. aeruginosa* colonies were counted after overnight incubation at 37 °C.

### Long-term gene circuit functionality

Ecc engineered with a GFP sensor circuit (Plasmid #6, Table S4) were used to assess long-term circuit activity. After 3 months of confinement, ILMs were induced with 100 μM 3OC6-HSL (Sigma-Aldrich), and GFP expression was monitored by time-lapse imaging. Additionally, after 1 or 6 months of *in vitro* confinement, or 7 days of *in vivo* confinement, ILMs were manually fragmented with a sterile razor blade and plated on agar. Individual colonies were picked, cultured, and induced with 100 μM 3OC6-HSL (Sigma-Aldrich). OD_600_ and GFP fluorescence were recorded every 20 minutes using a plate reader.

### Statistical analysis

Statistical analyses were performed using GraphPad Prism (v8). Tests, including Student’s *t*-test, one-way ANOVA, two-way ANOVA, linear regression, and nonlinear regression, were selected according to experimental design. The statistical test used for each figure is noted in the corresponding legend. Data are presented as mean ± SEM unless otherwise indicated. Statistical significance was defined as *P* < 0.05. When data were approximately normally distributed, comparisons were made using Student’s *t*-test (two groups), oneway ANOVA (single variable), or two-way ANOVA (two variables). Mice were randomized into groups before experiments.

