## Supplementary Materials for "Implantable Living Materials Autonomously Deliver Therapeutics from Contained Engineered Bacteria"

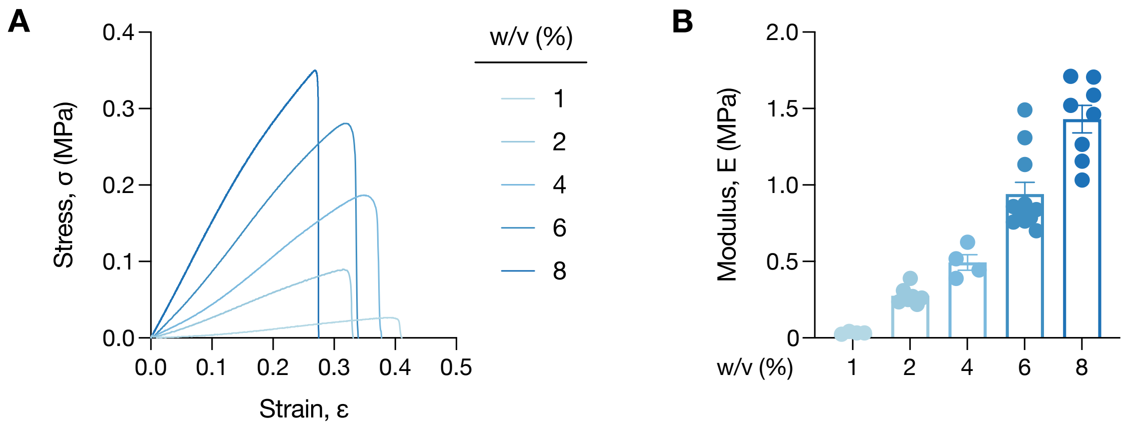


**Fig. S1 | Mechanical characterization of agarose hydrogels.**
(A) Representative stress–strain curves of UltraPure™ agarose hydrogels at different concentrations under uniaxial tension. (B) Elastic modulus of hydrogels shown in (A). Data are mean ± SEM; all replicates are shown.

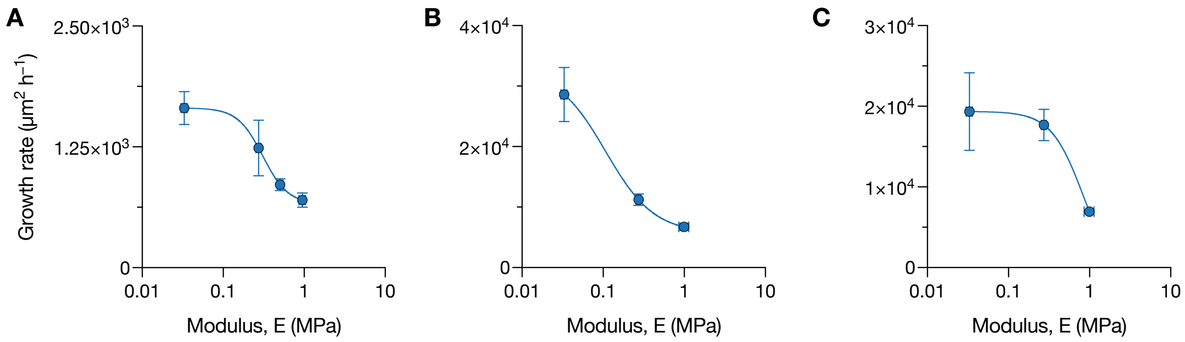


**Fig. S2 | Growth rate of three bacterial strains as a function of agarose stiffness.**
Growth rate of (A) *Escherichia coli* ClearColi™ BL21DE3 (Ecc), (B) *E. coli* Nissle 1917, and (C) *Bacillus subtilis* embedded in UltraPure™ agarose hydrogels. Data are mean ± SEM for n ≥ 2 biological replicates.

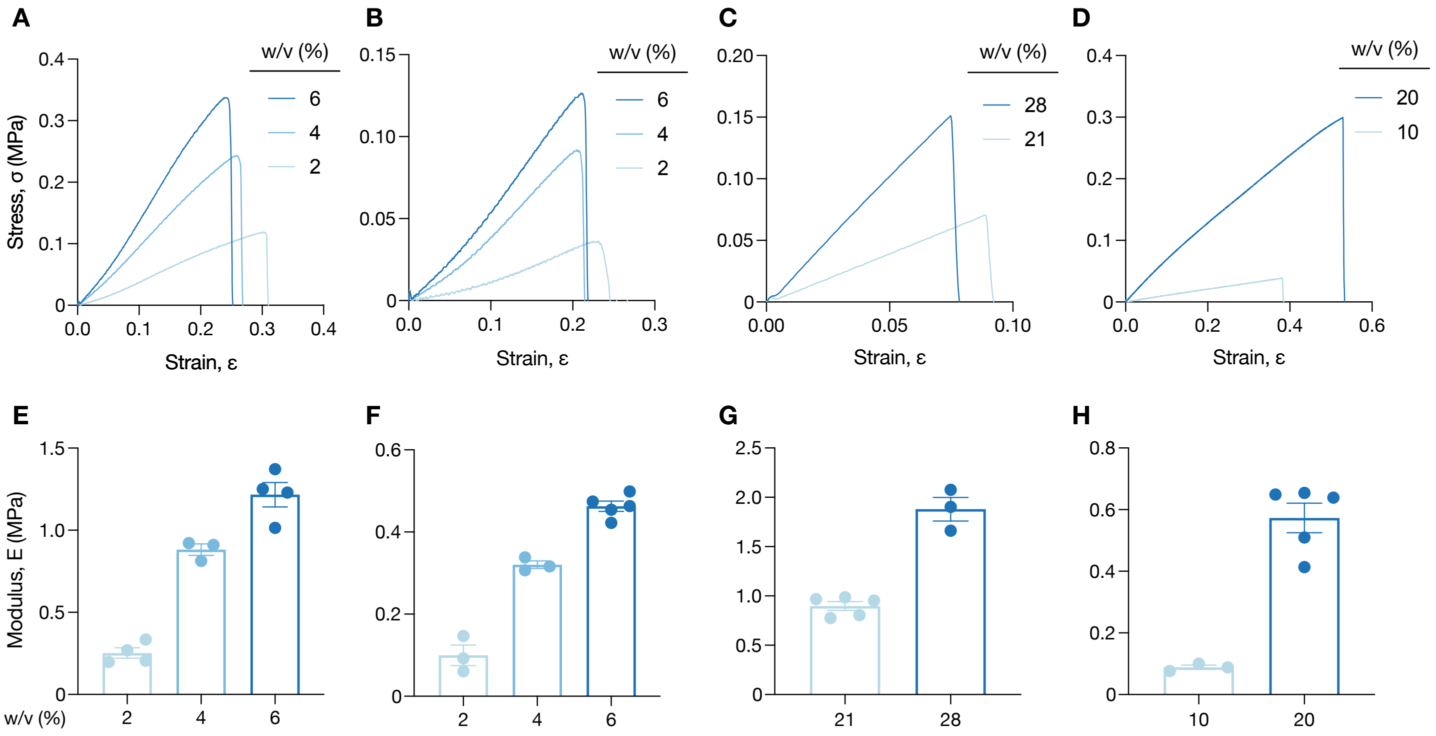


**Fig. S3 | Mechanical characterization of Certified Molecular Biology Agarose, Certified Low Range Ultra Agarose, PEGDA, and PEGDAAm hydrogels.**

Representative stress–strain curves of (A) Certified Low Range Ultra Agarose, (B) Certified Molecular Biology Agarose, (C) PEGDA, and (D) PEGDAAm hydrogels at different concentrations under uniaxial tension. (E–H) Elastic modulus of hydrogels shown in (A–D). Data are mean ± SEM; all replicates are shown.

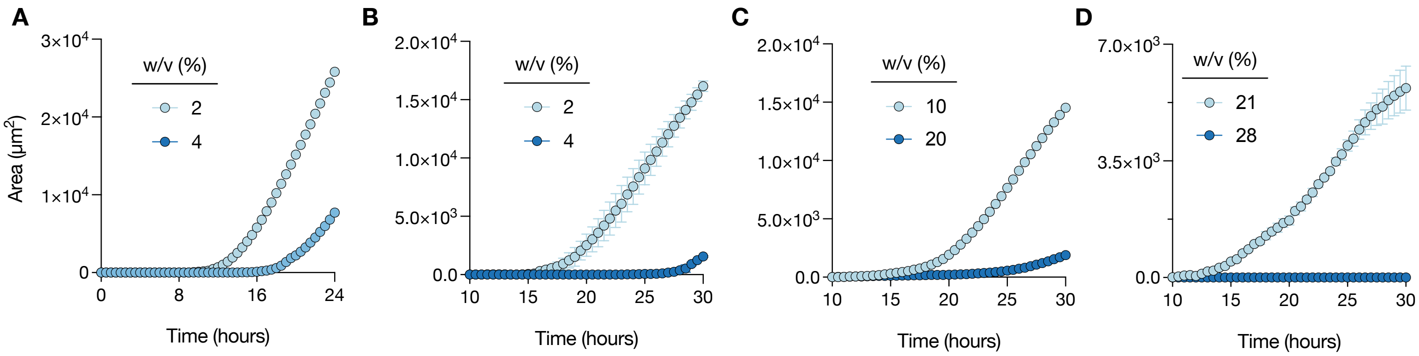


**Fig. S4 | Growth of encapsulated bacteria in agarose and PEG hydrogels of varying stiffness.** Quantification of bacterial colony area over time in (A) Certified Molecular Biology Agarose, (B) Certified Low Range Ultra Agarose, (C) PEGDA, and (D) PEGDAAm hydrogels at different molecular weights and concentrations. Ecc was embedded in each matrix. Data are mean ± SEM for n ≥ 2 biological replicates.

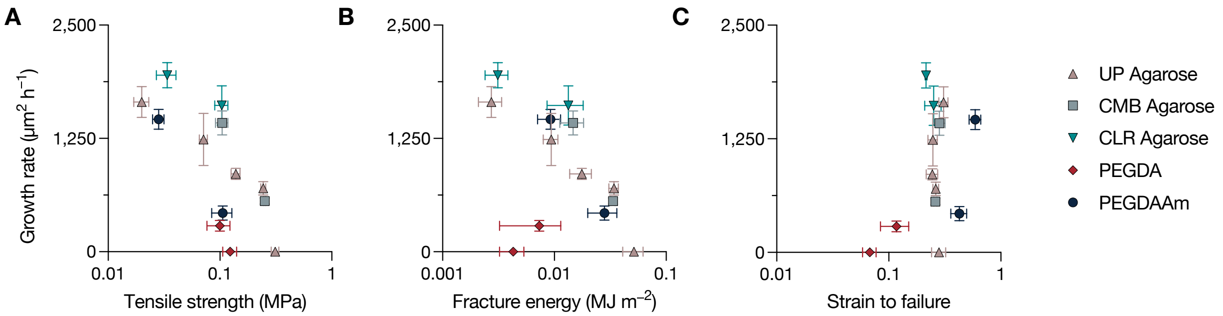


**Fig. S5 | Growth rate of embedded bacteria as a function of hydrogel mechanical properties.**
Growth rate of Ecc embedded in various hydrogel types as a function of (A) tensile strength, (B) work of fracture, and (C) fracture strain. Data are mean ± SEM for n ≥ 3 biological replicates.

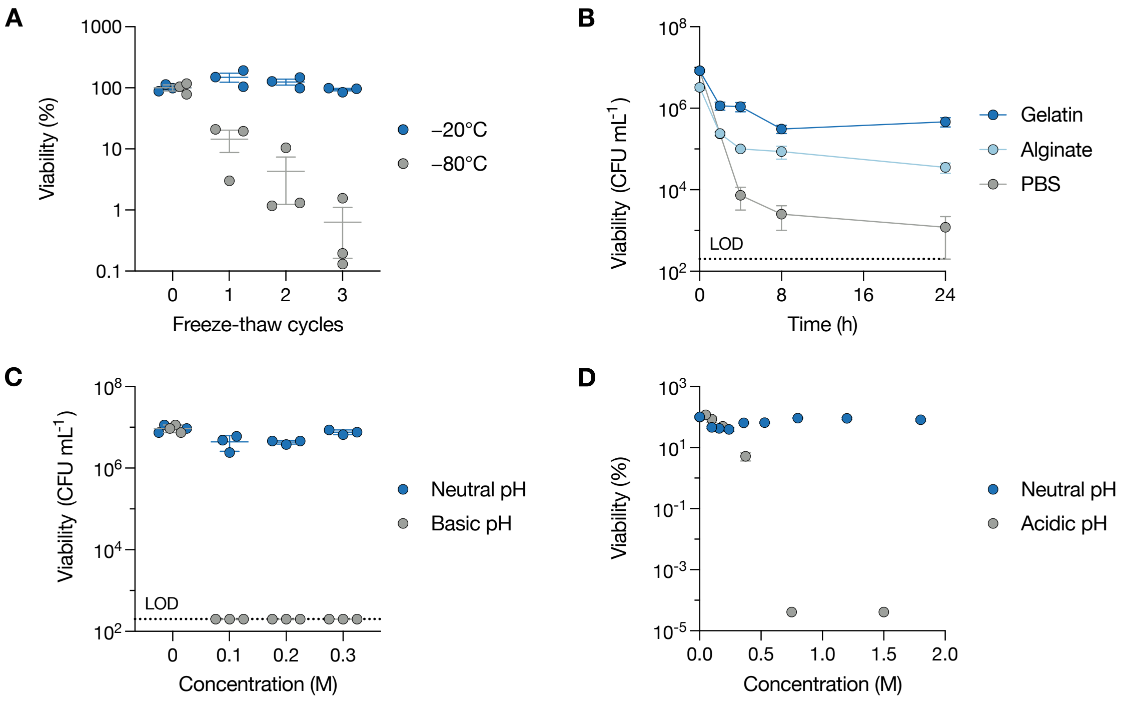


**Fig. S6 | Viability of encapsulated bacteria following ILM fabrication steps.**

(A) Viability of Ecc after repeated freeze–thaw cycles at −20 °C and −80 °C. (B) Viability of Ecc during dry annealing at 37 °C with or without incorporation of alginate (5% w/v) or gelatin (10% w/v). (C, D) Viability of Ecc after exposure to (C) sodium phytate or (D) sodium citrate salt solutions at different concentrations. Data are mean ± SEM for n = 3 biological replicates. Limit of detection (LOD) = $2 \times 10^2$ CFU ml$^{-1}$.

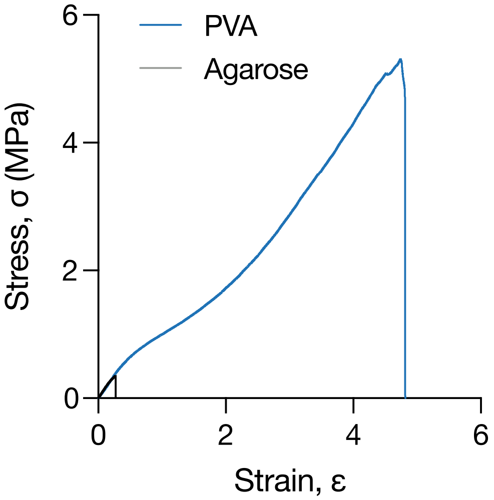


**Fig. S7 | Mechanical comparison of engineered PVA and agarose hydrogels with matched stiffness.** Representative stress–strain curves of engineered PVA hydrogels (Table S3, v) and UltraPure™ agarose hydrogels (8% w/v) under uniaxial tension.

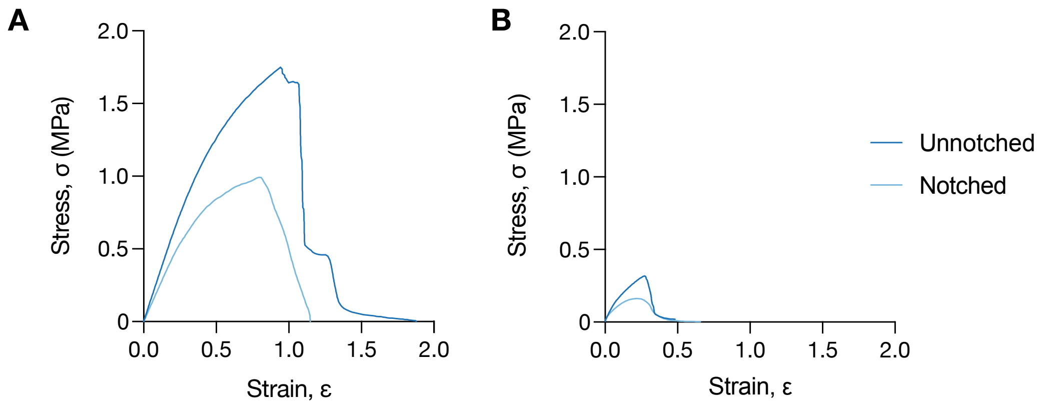


**Fig. S8 | Pure-shear tests of PVA and agarose hydrogels with matched elastic modulus.**
Representative stress–strain curves of (A) PVA hydrogels (Table S3, v) and (B) agarose hydrogels (8% w/v), both notched and unnotched, subjected to pure-shear tests.

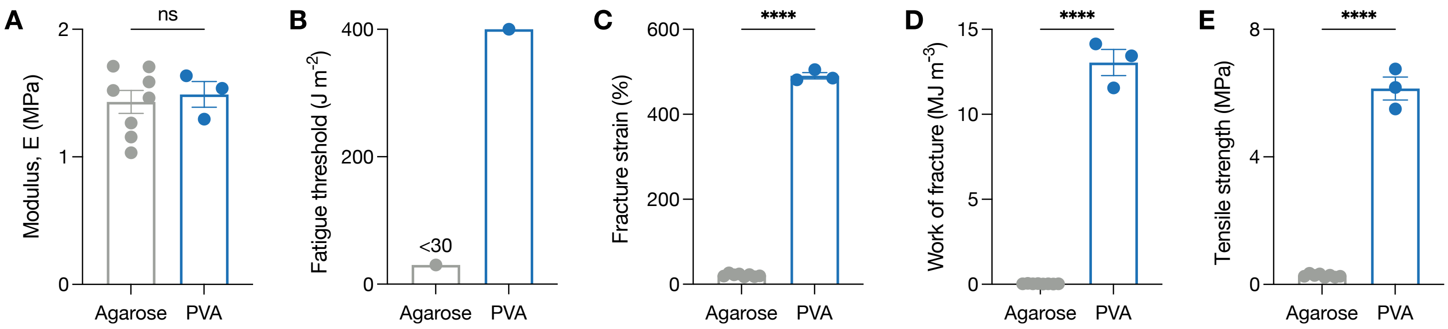


**Fig. S9 | Mechanical properties of engineered PVA and agarose hydrogels with matched elastic modulus.** Comparison of (A) stiffness, (B) fatigue threshold, (C) fracture strain, (D) work of fracture, and (E) tensile strength between agarose hydrogels (8% w/v) and engineered PVA hydrogels (Table S3, v). The fatigue threshold for agarose is roughly estimated due to its brittleness. Data are mean ± SEM. Statistical analyses were performed using two-sided unpaired t-tests (****$P < 0.0001$; ns, $P > 0.05$).

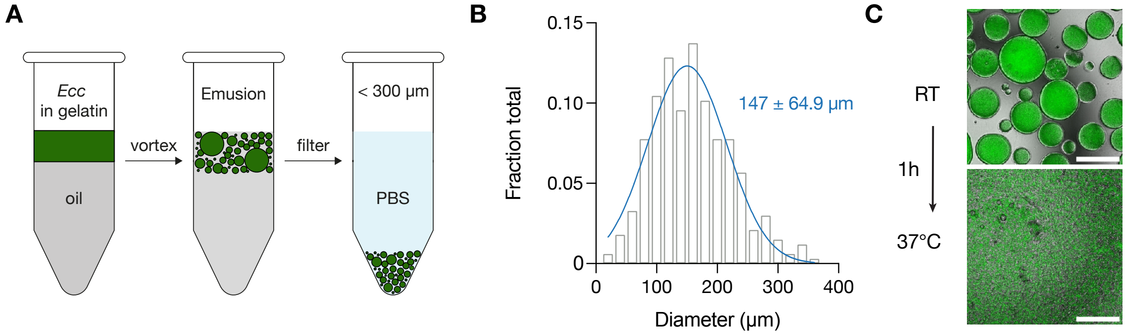


**Fig. S10 | Fabrication, size distribution, and thermoresponsiveness of bacteria-laden gelatin microgels.** (A) Fabrication steps of gelatin microgels. Ecc were suspended in 30% (w/v) gelatin, emulsified in oil by vortexing, gelled, washed, and filtered through a 300 μm strainer. (B) Size distribution of filtered gelatin microgels. (C) Representative fluorescence microscopy images showing the thermoresponsive behavior of gelatin microgels. Top: microgels at room temperature. Bottom: melted microgels at physiological temperature after 1 hour incubation. Scale bars, 200 μm.

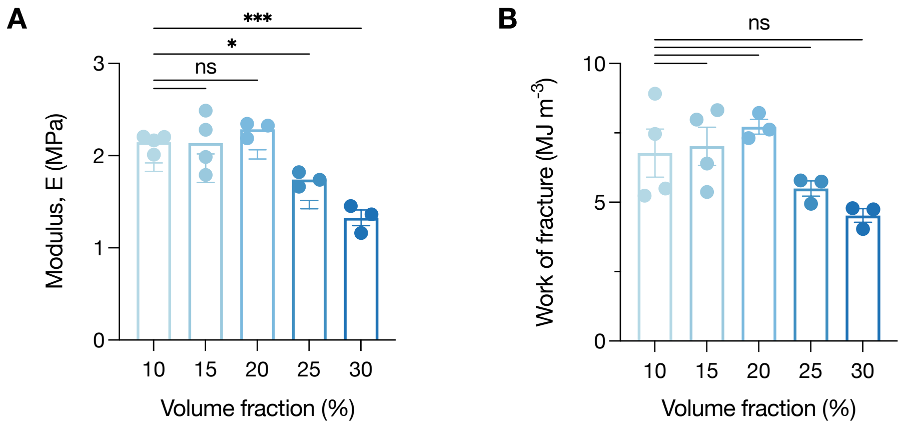


**Fig. S11 | Mechanical properties of engineered PVA hydrogels with varying microgel concentrations.** (A) Elastic modulus and (B) work of fracture of a PVA hydrogel containing different microgel volume fractions. Data are mean ± SEM. Statistical analyses were performed using one-way ANOVA with Dunnett's multiple comparisons test (***P = 0.0001; *P = 0.0451; ns, P > 0.05).

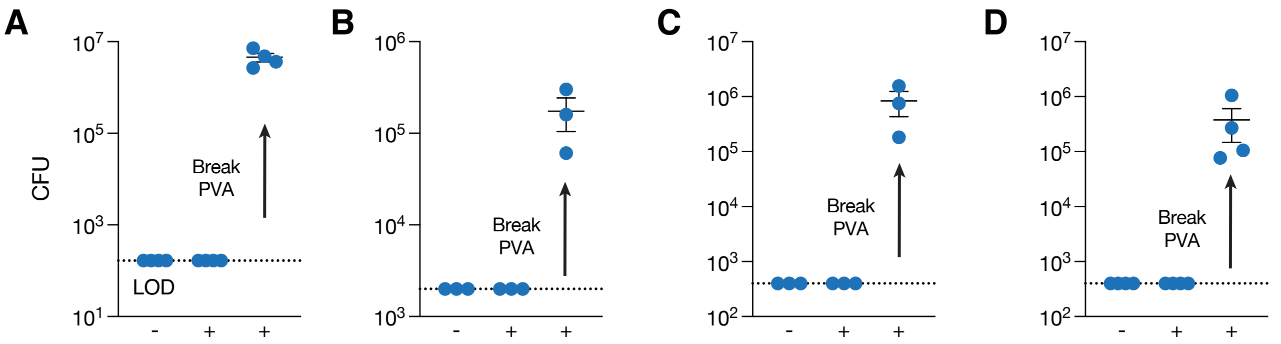


**Fig. S12 | Viability of encapsulated bacteria in ILMs after mechanical stress tests.**
(A–D) Viable colonies recovered from outside the PVA hydrogel after (A) compression, (B) tension, (C) tension with a notch, and (D) cyclic tensile loading. Notches in (C) and (D) simulated material defects. Bacterial counts were measured before (−) and after (+) loading. Samples were manually fractured to assess Ecc viability within ILMs after loading. Data are mean ± SEM. LOD = 166 (A), 2000 (B), 400 (C and D) CFU.

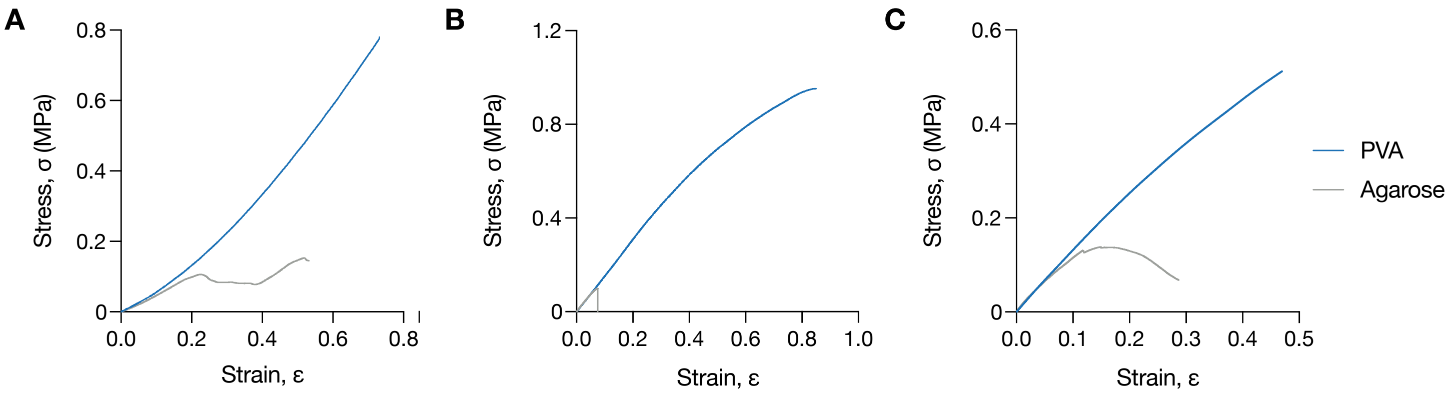


**Fig. S13 | Mechanical testing of PVA and agarose hydrogels.**
Representative stress–strain curves of PVA and agarose hydrogels under (A) compression, (B) uniaxial tension, and (C) pure shear.

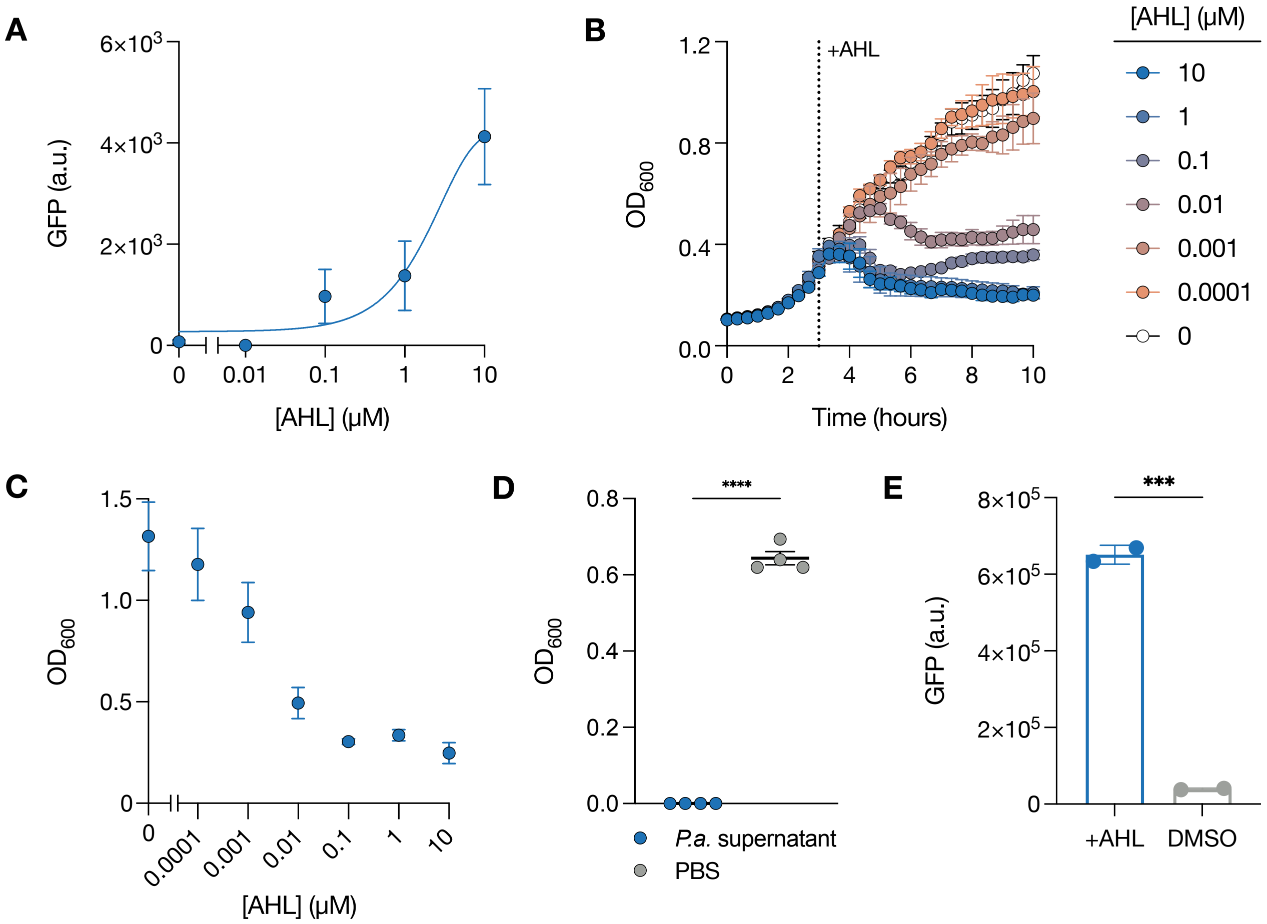


**Fig. S14 | Characterization of engineered Ecc with quorum-sensing circuits.**
(A) GFP expression of Ecc carrying the sensor circuit after induction with AHL at increasing concentrations. (B) Growth curves of Ecc carrying sensor and lysis circuits following induction with AHL. (C) $OD_{600}$ values of Ecc carrying sensor and lysis circuits after 10 hours of AHL exposure. (D) $OD_{600}$ values of Ecc carrying sensor and lysis circuits after incubation with 50% spent *Pseudomonas aeruginosa* medium (ATCC 10145) for 10 hours. (E) GFP release from Ecc carrying sensor and lysis circuits following induction with AHL for 3 hours. Data are mean ± SEM for $n \geq 3$ replicates. Statistical analyses were performed using two-sided unpaired t-tests (***$P = 0.0001$; ****$P < 0.0001$).

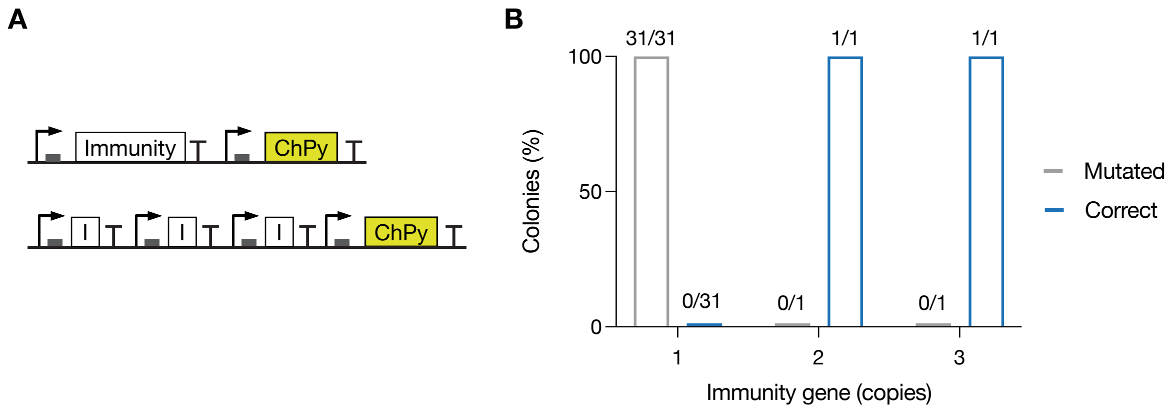


**Fig. S15 | Therapeutic payload circuit design in Ecc.**
(A) Schematic of engineered circuits in Ecc programmed to constitutively express a synthetic anti–*Pseudomonas aeruginosa* protein (ChPy) with either (top) a single immunity protein gene (I) or (bottom) multiple tandem immunity genes. (B) Summary of cloning outcomes after transforming the circuits shown in (A) into *E. coli* DH5a.

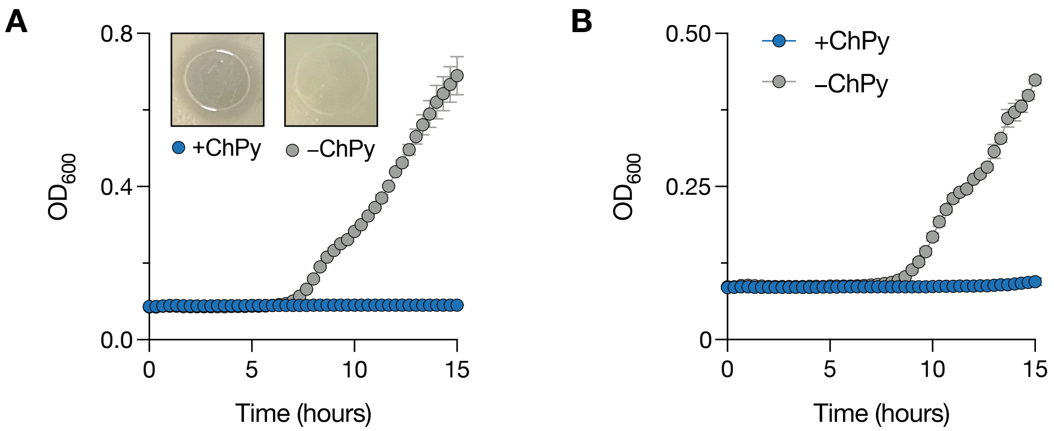


**Fig. S16 | Growth of *P. aeruginosa* strains in the presence of sonicated Ecc lysates.**
Growth curves of *P. aeruginosa* (A) BAA-47 and (B) Xen41 cultured with sonicated lysates from ChPy- or GFP-expressing Ecc. Insets in (A) show representative images of *P. aeruginosa* lawn regions spotted with the indicated lysates. Data are mean ± SEM for n = 4 biological replicates.

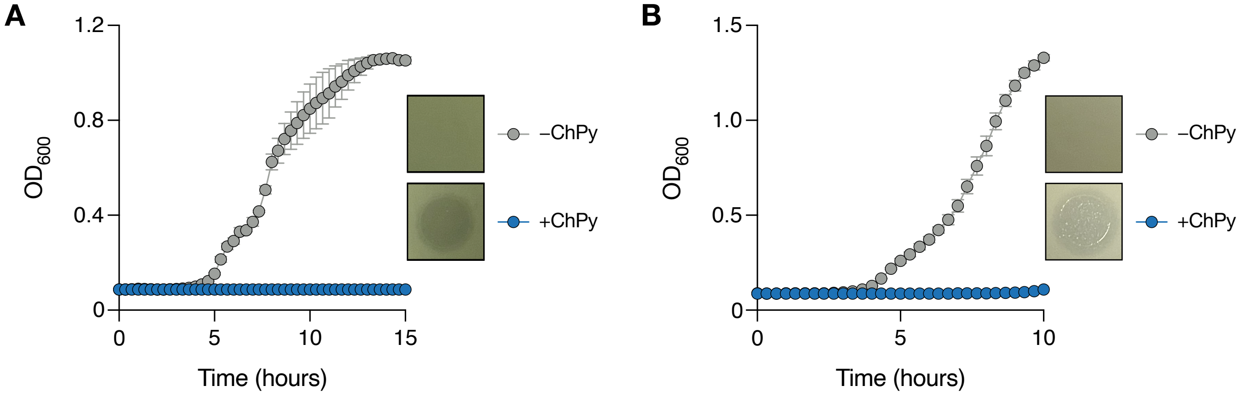


**Fig. S17 | Growth of *P. aeruginosa* in the presence of lysates from engineered Ecc.**
Growth curves of *P. aeruginosa* (A) ATCC 15692 and (B) BAA-47 cultured with ChPy-containing lysates. The lysates were prepared by exposing Ecc harboring the sensor–lysis–payload system to AHL for 3 hours. Insets show representative images of *P. aeruginosa* lawn regions spotted with lysates with or without ChPy lysates. Data are mean ± SEM for n = 3 biological replicates.

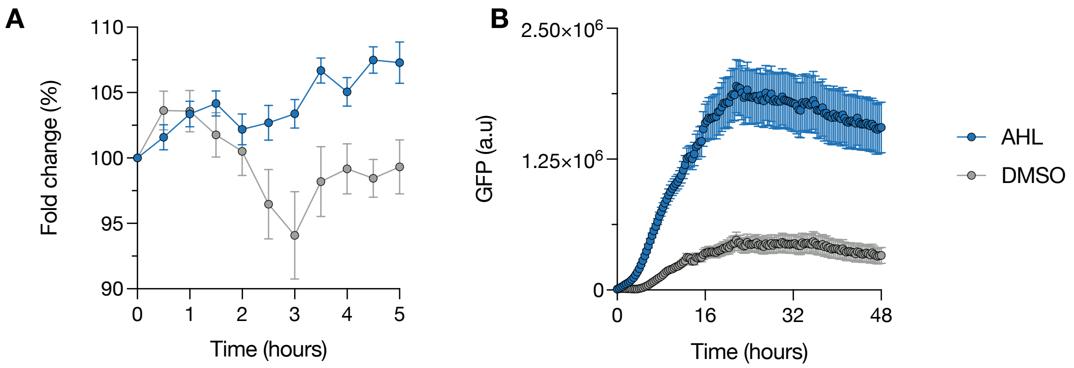


**Fig. S18 | Synthetic gene circuit activity after long-term bacterial encapsulation.**
(A) GFP expression of encapsulated Ecc upon induction with AHL after 3 months within ILMs. Data are mean ± SEM for n ≥ 3 biological replicates. (B) GFP expression of planktonic Ecc recovered from PVA hydrogel upon induction with AHL following 6 months of encapsulation. Data are mean ± SEM for n ≥ 3 biological replicates.

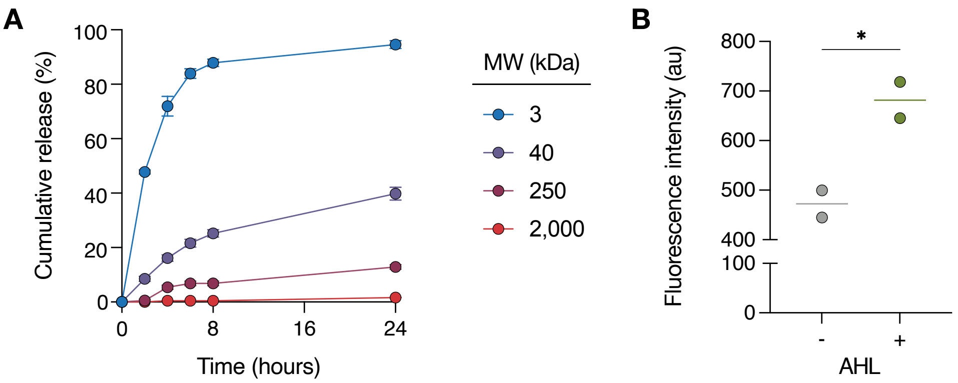


**Fig. S19 | Release of payloads from engineered PVA hydrogels.**

(A) Cumulative release profiles of fluorescein isothiocyanate–dextran (FITC–dextran, 3–2000 kDa) from engineered PVA hydrogels. Data are mean ± SEM for n = 3 replicates. (B) Payload release from ILMs. GFP expressed and released by Ecc within ILMs was detected in the surrounding medium upon induction. Data are mean ± SEM; all replicates are shown. *$P < 0.05$; unpaired *t*-test.

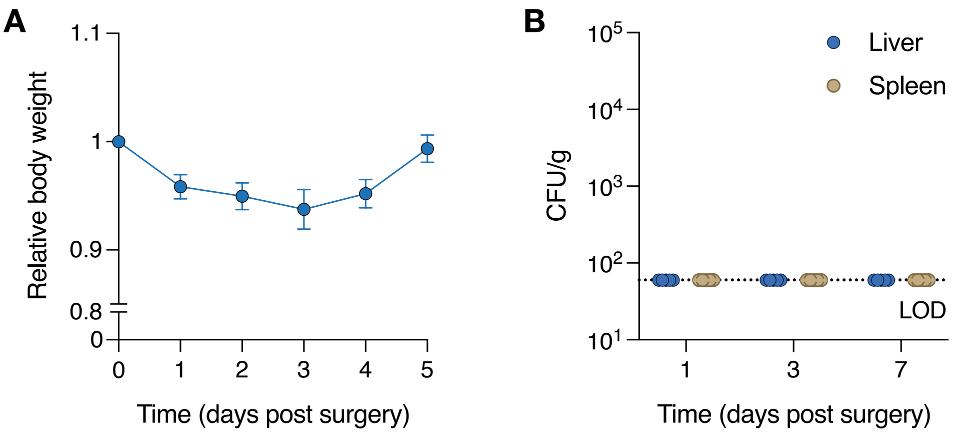


**Fig. S20 | Body weight and bacterial biodistribution in mice after ILM implantation.**
(A) Relative body weight of C57BL/6 mice following ILM and prosthetic joint implantation. (B) Ecc levels in the liver and spleen after ILM implantation. LOD = 60 CFU $g^{-1}$.

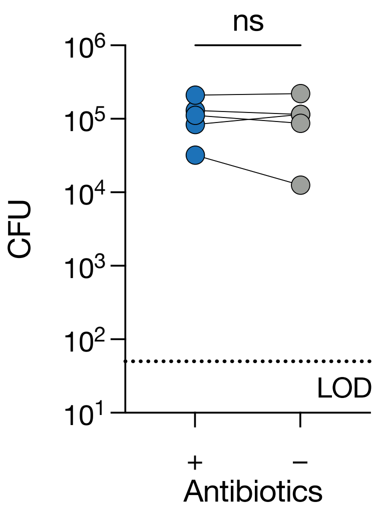


**Fig. S21 | Plasmid retention in encapsulated bacteria after *in vivo* implantation.**
Engineered Ecc CFU recovered from fragmented ILMs 7 days after implantation, plated with (+) or without (−) antibiotic selection. Data are from n = 5 biological replicates. Statistical analysis was performed using a two-sided paired t-test (ns, $P > 0.05$). LOD = 50 CFU $ml^{-1}$.

| Material | Bacteria | Application | Media | Containment Duration | Reference |
|---|---|---|---|---|---|
| Chitosan | *E. coli* | Subcutaneous implantation | Tissue simulation fluid | 12 hours* | Han et al., 2022 |
| Methacrylated hyarulonic acid | *E. coli, L. reuteri* | Cutaneous wounds | MRS | 12 hours | Ming et al., 2021 |
| Alginate | *S. elongatus* | Cutaneous wounds | BG11 | 5 days | Li et al., 2023 |
| Alginate + Polyacrylamide + Elastomer | *E. coli* | Wearable | LB | 24 hours | Liu et al., 2017 |
| Alginate | *E. coli* | Wearable | LB | 12 hours* | Allen et al., 2024 |
| PVA vinyl sulfone | *C. glutamicum* | Contact lenses | BHI | 21 days | Puertas-Bartolomé et al., 2024 |
| Elastin-like polypeptides | *E. coli* | Ingestion | LB | 2 days | Dai et al., 2021 |
| Hyarulonic acid + PEGDA + Mesoporous silica | *E. coli* | Environmental monitoring | LB | 3.5 days | Zhao et al., 2021 |
| Alginate methacrylate | *E. coli* | Environmental monitoring | LB | 4 hours* | Li et al., 2017 |
| Alginate + Polyacrylamide | *E. coli* | Environmental monitoring | LB | 3 days | Tang et al., 2021 |
| PEG diacrylate | *P. aeruginosa* | Environmental monitoring | LB | 23 hours | Steinacher et al., 2021 |
| Alginate | *E. coli* | Environmental monitoring | LB | 2 days* | Peng et al., 2023 |
| Pluronic F127 dimethacrylate | *P. aeruginosa* | Environmental treatment | Water | 5 days | Mayilsamy et al., 2025 |
| Pluronic F127 acrylate | *E. coli* | Biosensing | LB | 15 days* | Bhusari et al., 2023 |
| Agarose, Polyacrylamide | *E. coli* | Manufacturing | LB | 2 days | Sankaran et al., 2019 |
| Alginate | *L. plantarum* | Manufacturing | LB | 14 days | Tadimarri et al., 2025 |
| Chitosan | *E. coli* | Manufacturing | LB | 24 hours | Dai et al., 2019 |
| Alginate + Poly-L-lysine | *E. coli* | Coculture | M9 | 20 hours | Moya-Ramírez et al., 2022 |
| Alginate | *E. coli* | Coculture | LB | 10 days | Jeong et al., 2023 |

| | | | | | |
|---|---|---|---|---|---|
| Gelatin methacryloyl | *E. coli* | Coculture | LB | 24 hours* | Ou et al., 2023 |
| Pluronic F127-bisure-thane methacrylate | *E. coli* | Coculture | LB | 21 hours* | Johnston et al., 2020 |
| Alginate + poly-L-ly-sine | *E. coli, B. subtilis, P. putida, K. marxianus, Y. lipolytica* | Coculture | M9, LB, GPY | 2 days | Amaro-Cruz et al., 2025 |
| Chitosan | *E. coli, C. glutamicum, S. elongatus* | Coculture | M9 | 24 hours | Wang et al., 2022 |
| Gelatin methacryloyl | *E. coli* | Data storage | LB | 6 hours* | Luo et al., 2025 |

Table S1: Reported containment time of bacteria within hydrogel scaffolds. * indicates time to eventual bacteria leakage.

| Material | Source | Condition | Modulus (MPa) | Work of Fracture (J/m3) |
|---|---|---|---|---|
| Agarose | Experimental data | 1 wv% Agarose | 0.03 | 2.7 |
| | | 2 wv% Agarose | 0.27 | 9.3 |
| | | 4 wv% Agarose | 0.51 | 17.4 |
| | | 6 wv% Agarose | 0.93 | 34.1 |
| | | 8 wv% Agarose | 1.43 | 32.6 |
| PEG diacry-late | Experimental data | 14 wv% PEGDA | 0.16 | 3.1 |
| | | 21 wv% PEGDA | 0.93 | 3.2 |
| | | 28 wv% PEGDA | 1.88 | 4.3 |
| Pluronic F127 acry-late | Experimental data | 30 wv% F127, 25 wt% acry-lated F127 | 0.02 | 7.8 |
| | | 30 wv% F127, 50 wt% acry-lated F127 | 0.06 | 32.9 |
| | | 30 wv% F127, 75 wt% acry-lated F127 | 0.19 | 67.1 |
| | | 30 wv% F127, 100 wt% acry-lated F127 | 0.42 | 66.5 |
| Alginate | Sun et al., 2012 | 3 wt% Alginate, 10 wt% $CaSO_4$/Alginate | ~0.02 | ~3 |
| | | 3 wt% Alginate, 13.3 wt% $CaSO_4$/Alginate | ~0.05 | ~4 |
| | | 3 wt% Alginate, 16.6 wt% $CaSO_4$/Alginate | ~0.10 | ~6 |
| | | 3 wt% Alginate, 19.9 wt% $CaSO_4$/Alginate | ~0.14 | ~8 |
| | | 3 wt% Alginate, 23.3 wt% $CaSO_4$/Alginate | ~0.15 | ~10 |
| | | 3 wt% Alginate, 26.6 wt% $CaSO_4$/Alginate | ~0.17 | ~8 |
| Polyacryla-mide | Sun et al., 2012 | 13.6 wt% Acrylamide, 0.015 wt% MBAA/Acrylamide | ~0.007 | ~60 |
| | | 13.6 wt% Acrylamide, 0.031 wt% MBAA/Acrylamide | ~0.013 | ~100 |
| | | 13.6 wt% Acrylamide, 0.038 wt% MBAA/Acrylamide | ~0.014 | ~100 |
| | | 13.6 wt% Acrylamide, 0.046 wt% MBAA/Acrylamide | ~0.016 | ~100 |
| | | 13.6 wt% Acrylamide, 0.062 wt% MBAA/Acrylamide | ~0.016 | ~80 |
| | | 13.6 wt% Acrylamide, 0.108 wt% MBAA/Acrylamide | ~0.016 | ~50 |
| Alginate + Polyacryla-mide | Sun et al., 2012 | 66.67 wt% Acrylamide/(Algi-nate+Acrylamide) | ~0.1 | ~400 |
| | | 75 wt% Acrylamide/(Algi-nate+Acrylamide) | ~0.1 | ~1000 |
| | | 80 wt% Acrylamide/(Algi-nate+Acrylamide) | ~0.08 | ~1000 |
| | | 85.71 wt% Acrylamide/(Algi-nate+Acrylamide) | ~0.06 | ~2200 |
| | | 88.89 wt% Acrylamide/(Algi-nate+Acrylamide) | ~0.05 | ~2400 |

| | | | | |
|---|---|---|---|---|
| | | 92.31 wt% Acrylamide/(Alginate+Acrylamide) | ~0.02 | ~600 |
| | | 94.12 wt% Acrylamide/(Alginate+Acrylamide) | ~0.01 | ~300 |
| Gelatin methacryloyl | Shie et al., 2020 | 5 wv% Gelatin methacryloyl | ~0.05 | ~4 |
| | | 10 wv% Gelatin methacryloyl | ~0.08 | ~14 |
| | | 15 wv% Gelatin methacryloyl | ~0.14 | ~23 |
| Chitosan | Sampath et al., 2017 | 2 wv% Chitosan | ~0.01 | ~2 |

Table S2: Mechanical properties of hydrogels previously reported to encapsulate therapeutic *E. coli*

| | Molecular weight (kDa) | Concentration (w/v %) | Freeze/thaw cycle (number) | Freeze/thaw temperature (°C) | Freeze/thaw time (hour) | Annealing time (hours) | Salt type | Concentration (M) |
|---|---|---|---|---|---|---|---|---|
| a | 146-186 | 10 | 3 | -20 | 6 | - | Sulfate | 1.8 |
| b | 30-70 | 15 | 3 | -20 | 1 | - | Sulfate | 1.8 |
| c | 146-186 | 10 | 1 | -80 | 1 | - | Sulfate | 1.8 |
| d | 146-186 | 10 | 1 | -20 | 6 | - | Sulfate | 1.8 |
| e | 146-186 | 10 | 1 | -20 | 1 | - | Sulfate | 1.8 |
| f | 89-98 | 15 | 3 | -20 | 6 | - | Sulfate | 1.8 |
| g | 89-98 | 15 | 1 | -20 | 6 | - | Sulfate | 1.8 |
| h | 89-98 | 15 | 1 | -80 | 1 | - | Sulfate | 1.8 |
| i | 89-98 | 15 | 1 | -20 | 1 | - | Sulfate | 1.8 |
| j | 89-98 | 30 | 3 | -20 | 1 | - | Sulfate | 1.8 |
| k | 89-98 | 30 | 1 | -80 | 1 | - | Sulfate | 1.8 |
| l | 89-98 | 30 | 1 | -20 | 6 | - | Sulfate | 1.8 |
| m | 89-98 | 30 | 1 | -20 | 1 | - | Sulfate | 1.8 |
| n | 89-98 | 30 | 1 | -20 | 1 | 2 | Sulfate | 1.8 |
| o | 89-98 | 30 | 1 | -20 | 1 | 2 | - | - |
| p | 89-98 | 30 | 1 | -20 | 1 | 0.5 | Sulfate | 1.8 |
| q | 89-98 | 30 | 1 | -20 | 1 | 2 | Sulfate | 1.8 |
| r | 89-98 | 30 | 1 | -20 | 1 | 2 | Sulfate | 0.1 |
| s | 89-98 | 30 | 1 | -20 | 1 | 1.5 | Sulfate | 1.8 |
| t | 89-98 | 30 | 1 | -20 | 1 | 2 | Sulfate | 1.8 |
| u | 89-98 | 30 | 1 | -20 | 1 | 3 | Sulfate | 1.8 |
| v | 89-98 | 30 | 1 | -20 | 1 | 2 | Citrate | 0.1 |
| w | 89-98 | 30 | 1 | -20 | 1 | 2 | Citrate | 1.8 |
| x | 89-98 | 30 | 1 | -20 | 1 | 2 | Citrate | 0.6 |
| y | 89-98 | 30 | 1 | -20 | 1 | 2 | Phytate | 0.05 |
| z | 89-98 | 30 | 1 | -20 | 1 | 2 | Sulfate | 0.3 |
| A | 89-98 | 30 | 1 | -20 | 1 | 2 | Sulfate | 0.6 |
| B | 89-98 | 30 | 1 | -20 | 1 | 2 | Phytate | 0.3 |
| C | 89-98 | 30 | 1 | -20 | 1 | 2 | Citrate | 0.3 |
| D | 89-98 | 30 | 1 | -20 | 1 | 2 | Phytate | 0.2 |
| E | 89-98 | 30 | 1 | -20 | 1 | 2 | Phytate | 0.1 |

Table S3: PVA fabrication conditions

| Plasmid # | Plasmid | Promoter | ORI | Relevant Features |
|---|---|---|---|---|
| 1 | Sensor GFP (pAJM.474) | pLacI + pLuxB | p15A | OC12 sensor, LuxR + pLuxB-YFP reporter |
| 2 | Sensor mKate | pLacI + pLuxB | p15A | OC12 sensor, LuxR + pLuxB-mKate reporter |
| 3 | Lysis | pLuxB | sc101m | AHL-inducible bacteriophage lysis gene, $\varphi$X174 E |
| 4 | Constitutive ChPy | BBa_J23101 | pUC | Chimeric pyocin gene + 3 immunity genes (constitutive expression) |
| 5 | Constitutive GFP | BBa_J23101 | pUC | Constitutive GFP expression |
| 6 | pTD103 GFP (no luxI) | pLuxI | ColE1 | AHL-inducible GFP expression |

Table S4: Main plasmids used in this study.

**Movie S1**
**Bacteria growth in a polymeric hydrogel.** *E. coli* grows within a 2% agarose hydrogel over time. Scale bar, 50 μm.

**Movie S2**
**ILM and agarose under compressive stress.** Hydrogels were subjected to compressive stress up to ~0.8 MPa. Playback speed, 5x.

**Movie S3**
**ILM and agarose under tensile stress.** Hydrogels were subjected to tensile stress up to ~1 MPa. Playback speed, 20x.

**Movie S4**
**ILM and agarose with pre-existing defects under tensile stress.** Notched hydrogels were subjected to tensile stress up to ~0.5 MPa. Playback speed, 10x.

**Movie S5**
**ILM and agarose with pre-existing defects under cyclic tensile stress.** Notched hydrogels were subjected to repeated cycles of 7.5% strain. Playback speed, 20x.

**Movie S6**
**Gene circuit dynamics in ILM.** Addition of AHL to the surrounding media triggers mKate expression and subsequent lysis of embedded *E. coli*. Scale bar, 50 μm.
